# Data-Driven Retrieval of Effective Point Spread Functions for Super-Resolution Optoacoustic Imaging

**DOI:** 10.64898/2026.01.26.701644

**Authors:** Hongtong Li, Haiyang Zhan, Rui Ma, Yijun Liu, Hongjing Cao, Tingkai Yan, Zheng You, Xose Luis Dean-Ben, Daniel Razansky, Fei Xing

## Abstract

Localization-based super-resolution techniques have revolutionized biomedical imaging by surpassing classical diffraction limits. However, their performance is fundamentally constrained by distortions of the point spread function (PSF) induced by the system and the sample, which are particularly prominent in the case of spatially under-sampling. Here we introduce a data-driven effective point spread function retrieval (DEPR) method that directly learns continuous, field-dependent system responses from experimental point source datasets. Through statistical aggregation of thousands of targets and iterative self-supervised refinement, DEPR captures spatially variant imaging characteristics *in situ* without prior assumptions or external calibrations. When integrated into the localization optoacoustic tomography (LOT) pipeline, DEPR achieves accurate sub-pixel localization despite under-sampled conditions, thus enhancing resolution while reducing computational burden. We demonstrate its efficacy by *in vivo* imaging of the murine brain microvasculature using microparticle contrast agents in the first (NIR-I) and second (NIR-II) near-infrared windows, achieving significant improvement in data utilization efficiency and substantial reduction of gridded artifacts compared to conventional approaches. The method resolves vascular structures with ∼41 μm separation across a ∼3 mm imaging depth range. This framework addresses fundamental challenges shared across diverse localization-based imaging modalities, offering a robust and generalizable strategy for high-precision imaging in complex biological systems.

## Introduction

Super-resolution imaging has transformed optical microscopy and other modalities, enabling visualization of microscopic structures with unprecedented detail^1,2^. By expanding the set of distinguishable features, these approaches enhance diagnostic precision and illuminate complex biological processes^3–5^. Among them, localization-based techniques, first demonstrated in optical nanoscopy, overcome the classical diffraction limit by temporally isolating and localizing individual blinking targets, enabling nanoscale visualization of subcellular dynamics^6–9^. Ultrasound imaging subsequently adopted a similar framework to achieve super-resolution angiography by localizing and tracking microbubbles^10,11^. More recently, optoacoustic (OA) imaging has matured into a versatile platform that combines optical contrast with ultrasonic resolution for functional and molecular imaging at depths beyond the reach of purely optical microscopy methods^12–16^. Building on this foundation, localization optoacoustic tomography (LOT) preserves the advantages of OA while surpassing the acoustic diffraction limit^17–22^. This is achieved by localizing flowing absorbers, thereby facilitating mapping the microvasculature and hemodynamics at high resolution. Crucially, OA further offers label-free readouts such as blood oxygenation^23,24^, positioning LOT as a unique multiparametric modality with broad relevance in the fields of oncology, neuroscience, metabolism, and other applications^25–30^.

High-quality LOT reconstructions, however, hinge on precise absorber localization. Conventional strategies fall into centroid-based and model-fitting categories. Centroid methods estimate position via center-of-gravity (CG) calculations and are computationally efficient, but they exhibit intrinsic bias when the point spread function (PSF) becomes narrow relative to the volumetric pixel size^31^. This pixelation effect introduces systematic, S-shaped localization errors that compromise sub-pixel accuracy. Model-fitting approaches improve accuracy by matching observed intensities to parametric PSF models (e.g., Gaussian)^32,33^, but they are computationally intensive and sensitive to model mismatch. In practice, aberrations from inhomogeneous system response and sample characteristics distort the PSF from its theoretical form^34,35^, introducing localization errors particularly problematic in volumetric imaging within sub-Nyquist under-sampled scenarios.

Several strategies have been attempted to address these limitations. Reducing pixel size ensures the PSF spans multiple volumetric pixels (also referred to as voxels), improving localization precision at the expense of significantly higher data volume, computational demands, and artifact susceptibility^36^. Since obtaining a single super-resolved image requires tens of thousands of frames, this approach becomes resource-intensive and difficult to scale. External PSF calibration through imaging an isolated point source serves as another alternative^37,38^, but this *ex vivo* procedure fails to capture other reconstruction inaccuracies present in biological specimens, such as variations in the speed of sound. Recent developments have explored reconstructing PSFs directly from experimental data^39,40^, but most require strong priors and produce discretized representations that inadequately account for pixelation effects, making them unsuitable for under-sampled conditions.

To more faithfully describe the image acquisition and rendering pipeline, the effective point spread function (ePSF) has emerged as a continuous, experimentally derived model that integrates physical, electronic, and digital effects^41,42^. Successfully implemented in telescopic and other imaging systems, ePSF-based localization demonstrates strong performance in mitigating systematic errors^43,44^. However, current approaches lack adaptability to dynamic conditions encountered *in situ*. As such, accurate modeling of *in situ* ePSFs from experimental data remains challenging, particularly in under-sampled scenarios where pixelation-induced distortions dominate, which continuously hinders efforts toward achieving robust, high-fidelity super-resolution LOT imaging of biological structures.

Here, we developed a data-driven effective point spread function retrieval (DEPR) method that reconstructs continuous, field-dependent ePSFs directly from blinking datasets acquired within biological specimens. Unlike conventional approaches relying on external calibration or predefined models, DEPR captures the combined effects of aberrations, pixelation, and sample-induced distortions *in situ*. Conceptually, DEPR functions as a self-supervised iterative framework that pools heterogeneous data, extracts stable features, and progressively refines the model. This further enables accurate sub-pixel localization even in spatially under-sampled conditions. With integration into LOT, the proposed DEPR-based strategy enhances resolution and imaging fidelity while reducing data and hardware demands. We further demonstrate its application *in vivo* for mapping the brain vascular structure of a mouse, highlighting its potential for scalable, high-precision imaging in complex biological systems.

## Results

### Fast super-resolution optoacoustic imaging with microparticles

We implemented LOT to visualize the cerebral microvasculature in mice with super-resolution. The basic principle and workflow are illustrated in Fig. 1a and Supplementary Fig. S1. After intravenous (i.v.) injection of microparticles, repetitive laser pulses excited OA signals from these as they circulate within the brain vasculature. Due to their size, the particles remained confined within the vascular lumen without extravasation, enabling angiographic contrast enhancement. Raw signals were recorded by a custom-designed spherical transducer array (Fig. 1b) and reconstructed into volumetric images using a GPU-accelerated filtered back-projection (FBP) algorithm (see methods for details). Conventional OA tomography revealed large cerebral vessels, albeit with resolution limited by acoustic diffraction (Fig. 1c). To separate moving particles from background vasculature signals, singular value decomposition (SVD) was applied to isolate dynamic components across image sequences (Fig. 1d). We further provide exemplary demonstration of the reconstructed OA tomography images under both fine-sampled (volumetric pixel size: 0.06 mm) and under-sampled (volumetric pixel size: 0.12 mm) conditions. Although OA tomography images can be theoretically reconstructed with arbitrarily small volumetric pixel sizes, this does not improve the intrinsic effective resolution but exponentially increases computational and memory demands, limiting practical scalability^36^. The final LOT image was rendered from localizing and accumulating individual particle positions over time, revealing microvascular details well beyond the native resolution of the system (Fig. 1e). Comparably, these structures remained unresolved in diffraction-limited OA tomography images. Imaging was conducted without inducing observable physiological changes, confirming the safety and compatibility of the method with *in vivo* experimentation.

**Figure 1.**
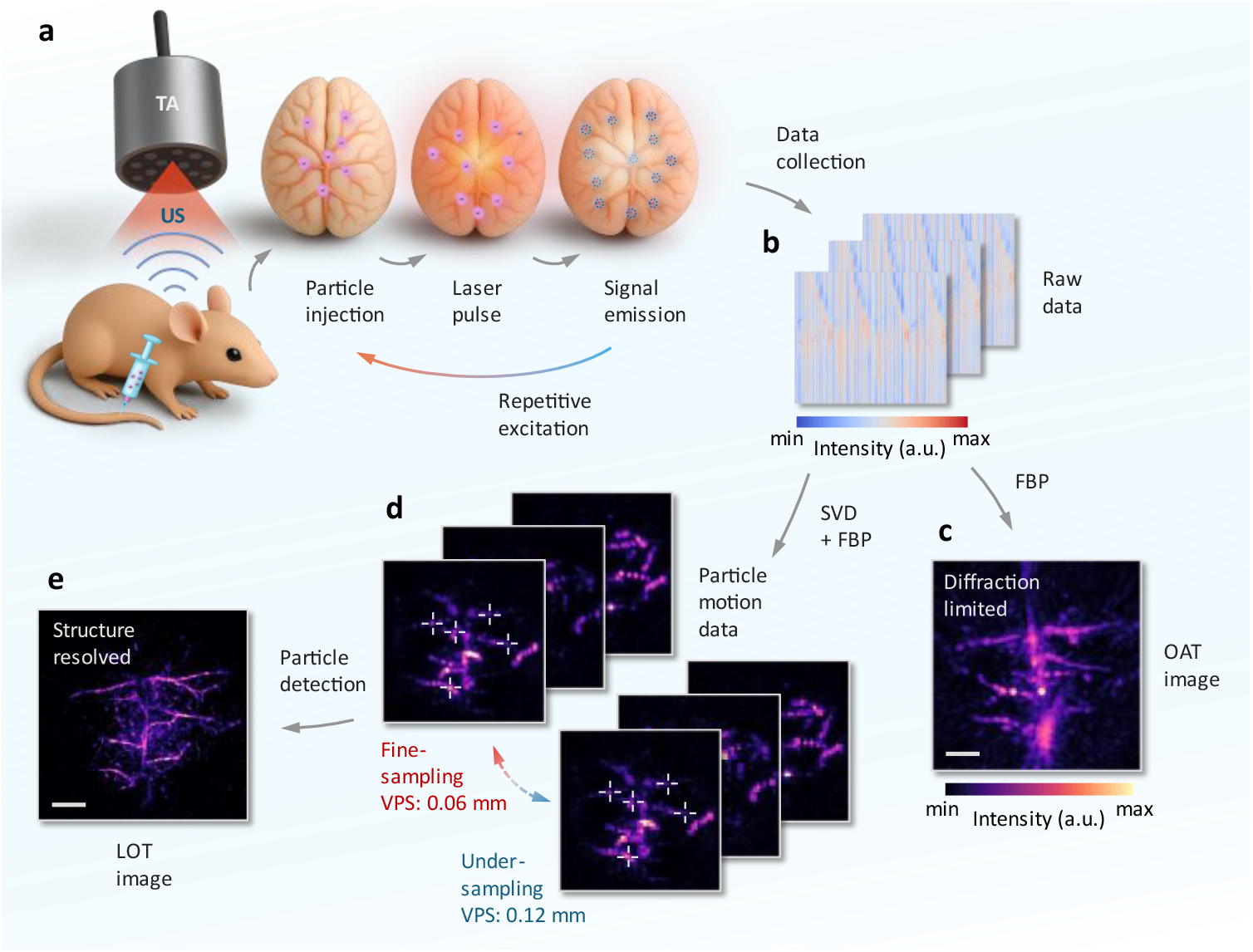
Localization optoacoustic tomography (LOT) for super-resolution vascular imaging in the mouse brain. **a**, Schematic of the LOT imaging process. Microparticles are intravenously injected and excited via pulsed laser illumination. Emitted ultrasound (US) signals are detected by a spherical transducer array (TA) with repeated excitation enabling dynamic tracking. **b**, Raw signals are acquired over time and reconstructed using an universal filtered back-projection (FBP) algorithm, which incorporates band-pass filtering and GPU-accelerated back-projection steps. **c**, Standard optoacoustic tomography (OAT) image showing major cerebral vasculature with resolving ability limited by acoustic diffraction. **d**, Individual microparticle trajectories are extracted from motion-resolved images using singular value decomposition (SVD). Comparisons between fine-sampled data with volumetric pixel size (VPS) of 0.06 mm and under-sampled data with VPS of 0.12 mm demonstrate distinct reconstruction outcomes. Each panel includes maximum intensity projections (MIPs) of 30-frame accumulations, providing clearer visualization of sampling differences. **e**, LOT image formed by accumulating localized positions across frames, resolving vascular structures beyond the diffraction limit. Scalebar – 1 mm, a.u. – arbitrary units.

### Data-driven retrieval of ePSFs for enhanced localization

The PSF defines the response to a point source. In practice, it often departs from ideal models (e.g. Gaussian) due to aberrations, defocus, and detector sampling. Pixelation is particularly detrimental, producing spatially varying, anisotropic, discretized responses that bias localization. The ePSF concept addresses this by combining the continuous physical response with discrete sampling effects, enabling accurate sub-pixel localization from pixelated data^42,45^. DEPR reconstructs high-resolution, field-dependent ePSFs directly from routine datasets without external calibration. Embedded in the LOT pipeline, DEPR aggregates randomly distributed absorbing point sources and iteratively learns the continuous, spatially variant system response. Unlike fixed, pixel-discrete PSF models, the resulting ePSFs are continuous and sub-pixel resolved, effectively recovering fine-sampled behavior from under-sampled measurements.

The workflow begins with conventional FBP reconstruction and dynamic background subtraction (Fig. 2a). The field of view (FOV) is partitioned into subregions within which the system response is assumed locally invariant. Regions of interest (ROIs) around detected microparticles are grouped according to their spatial location (Fig. 2b). To account for their sub-pixel displacements, each ROI is spatially aligned to its estimated centroid position and assigned to a discrete sub-pixel bin representing its fractional offset within the native volumetric pixel grid (Fig. 2c). In essence, this binning process converts randomly distributed target positions into a structured sampling of the underlying continuous PSF. The selection of sub-pixel bin resolution represents a critical trade-off between reconstruction accuracy and computational efficiency. Finer sampling improves precision but increases computational cost and reduces statistical robustness, whereas coarser sampling lowers accuracy. Optimal parameters should therefore be adapted to specific imaging conditions and sampling density (Supplementary Fig. S2). To construct the ePSF, pixel intensities from all aligned ROIs are mapped into their corresponding relative coordinates (Fig. 2d), and data from thousands of such observations are aggregated with statistical filtering to reject outliers. The accumulated measurements, spanning a dense range of sub-pixel positions, are then interpolated using cubic splines to generate smooth, continuous ePSFs for each subregion (Fig. 2e). Crucially, this approach achieves genuine resolution enhancement by recovering intra-pixel structure from aggregated experimental observations, in contrast to conventional interpolation that merely smooths discretized data.

**Figure 2.**
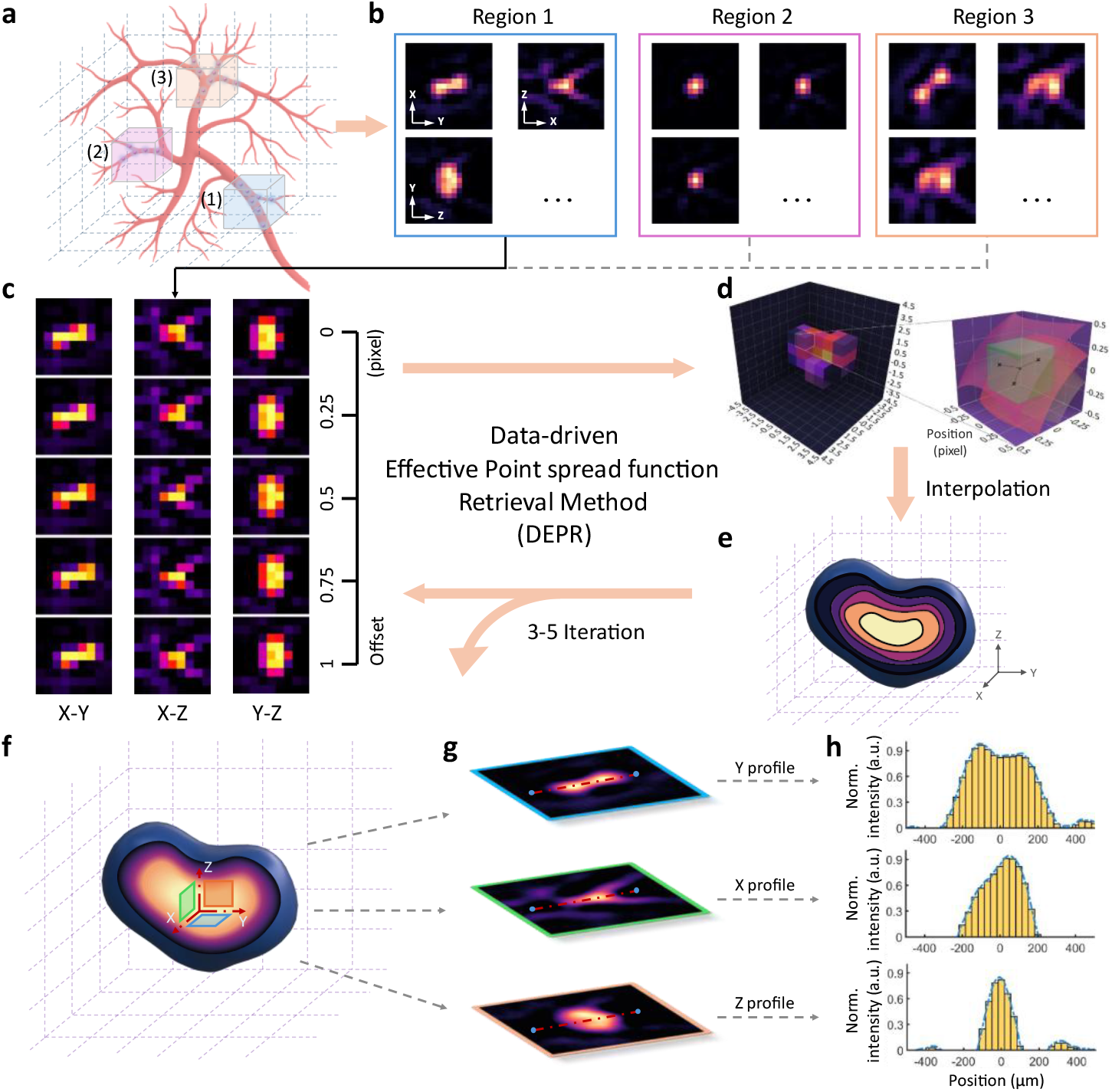
Workflow of data-driven effective point spread function retrieval (DEPR). **a**, Schematic illustration of dividing the imaging volume into spatial subregions to account for field-dependent variations. **b**, Representative raw particle regions of interest (ROIs) collected from different subregions, exhibiting distinct point spread function (PSF) characteristics. Maximum intensity projections (MIPs) along all three Cartesian directions (x,y,z) are shown. **c**, ROIs are further classified into sub-pixel bins according to estimated centroid offsets along x, y, and z axes. **d**, Aggregated PSF samples are aligned and assigned to their corresponding sub-pixel bins, producing pixelated PSF representations that preserve intra-pixel structure. **e**, Interpolation converts pixelated PSFs into smooth, continuous effective point spread functions (ePSFs). **f**, Iterative refinement progressively improves the reconstructed ePSFs. **g**, Final ePSFs are visualized as three orthogonal cross-sections, demonstrating smooth, high-resolution representations. **h**, Normalized intensity profiles along the x, y, and z axes corresponding to g.

An iterative self-supervised update loop then refines both absorber localization and model accuracy. Least-squares fitting (LSF) against the reconstructed ePSFs yields improved sub-pixel centroiding results, which are propagated back into the binning step. Analogous to iterative model training, each cycle improves the accuracy of both the data labels (microparticle positions) and the learned representation (ePSF model). Convergence typically occurs within a few iterations (Supplementary Fig. S3). The final output is a field-dependent library of high-resolution ePSFs that capture local system characteristics across the imaging volume (Fig. 2f). Notably, DEPR requires no prior PSF assumptions and generalizes seamlessly from one-to three-dimensional reconstruction. More detailed derivations are provided in the methods section. The ePSFs recovered by DEPR provide useful physical insights, and can serve to derive refined sub-pixel centroid estimates. Accumulating these localizations over time yields super-resolved angiograms that exceed the acoustic diffraction limit and remain robust for under-sampled conditions.

### Quantitative field-dependent variation and sub-pixel localization assessment

To assess field-dependent variations in system response, we first performed an external calibration using a sub-resolution microparticle (diameter ∼35 µm) embedded in an agar phantom. Considering the axial symmetry of the spherical array, the phantom was translated across a 12×12 spatial grid with 0.5 mm steps covering one quarter of the full FOV, and volumes were reconstructed with the standard FBP algorithm^36^. With implementation of our previously reported ePSF characterization workflow^44^, we compared a library of conventional discrete PSFs with continuous ePSFs across the tomographic volume. The results revealed substantial non-uniformity in system response, particularly toward the FOV periphery (Fig. 3a). Relative to conventional PSFs, the established ePSFs captured rich intra-pixel structure and anisotropy, avoiding shape distortions and yielding more informative lateral profiles (Fig. 3b). These findings expose a fundamental limitation of centroid-based localization in under-sampled conditions, where pixelation-induced bias dominates. Frequency-domain analysis further substantiated the spatial heterogeneity, with one-dimensional Fourier transforms of raw OA signals revealing position-specific spectral signatures across the field (Supplementary Fig. S4). We further quantified the variations by principal component analysis (PCA)-based projecting the calibrated full-field ePSF and computing its Euclidean distance from the central ePSF^46^. The resulting spatial map revealed systematic trends, including progressive broadening and increased ellipticity toward the edges of the FOV (Fig. 3c), thereby underscoring the necessity of field dependent modeling for high precision localization.

**Figure 3.**
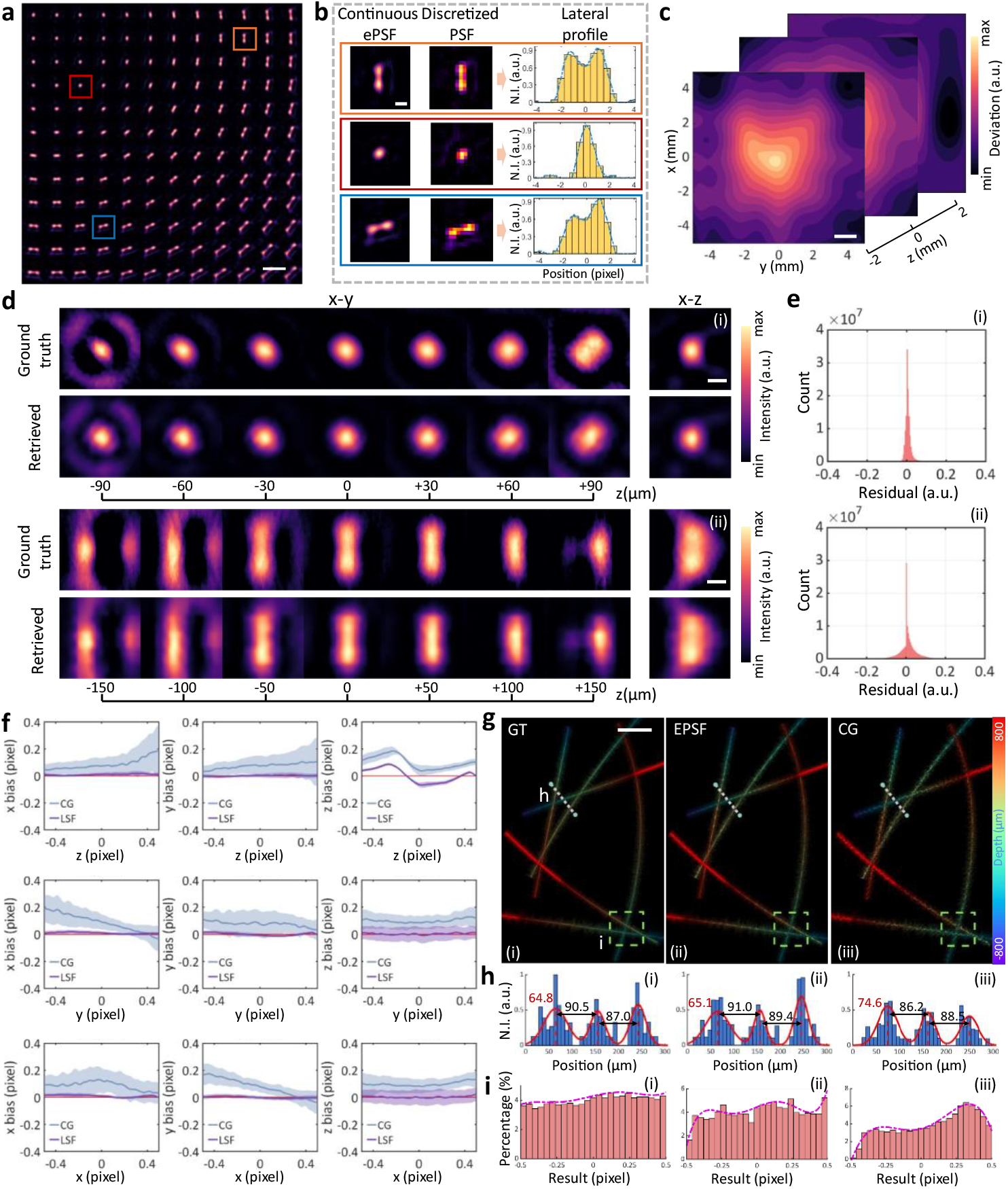
Validation of DEPR with experimental calibrations and simulations. **a**, External calibration with sub-resolution particles embedded in an agar phantom sampled on a 6 × 6 mm^2^ grid covering one quarter of the full field of view (FOV). Scalebar – 500 μm. **b**, Comparison between discretized PSFs and DEPR-reconstructed continuous ePSFs from selected subregions, with corresponding lateral profiles at right illustrating intra-pixel structure recovered by DEPR. Scalebar – 200 μm. **c**, Principal component analysis (PCA) of calibrated PSFs across the FOV, with spatial mapping of deviations from the central PSF revealing systematic broadening and ellipticity toward the periphery. Scalebar – 1 mm, a.u. – arbitrary units. **d**, Comparison of ground truth (GT) and DEPR-reconstructed ePSFs in x–y and x–z planes, showing close agreement in anisotropy and intensity falloff. Scalebar – 100 μm. **e**, Residual distributions after normalization demonstrate pixel-level fidelity, with errors largely confined within ±10%. **f**, Localization accuracy benchmark using simulated absorbers at controlled sub-pixel positions. DEPR-based least-squares fitting (LSF) consistently reduces systematic bias compared to center-of-gravity (CG) estimation, particularly in the axial direction. **g**, Structural reconstruction of synthetic microtubule networks. GT, CG, and DEPR-based LSF results are shown side by side, where the proposed approach preserves filament continuity and suppresses gridded artefacts. Scalebar – 200 μm. **h**, Enlarged regions and line profiles confirm sharper resolution, improved contrast, and better agreement with the GT. **i**, Sub-pixel localization histograms show markedly better agreement with reference distributions using DEPR, indicating reduced position-dependent bias.

To validate the retrieval of the imaging response, we generated synthetic volumetric datasets using ground truth PSFs sampled from experimental calibrations. We simulated randomly placed sub-pixel absorbers, generated noisy under-sampled observations from their ground truth PSFs, and processed them with DEPR using the same iterative pipeline as for real data. Fig. 3d shows side by side comparisons of ground truth and DEPR reconstructions in both x-y and x-z planes with close agreement in shape, anisotropy, and intensity falloff. Residual maps confirmed accurate recovery across central and peripheral regions, and pixel level differences after normalization were confined for the great majority to within ±10%, as summarized by the narrow residual histograms in Fig. 3e. Performance remained robust across a range of signal-to-noise ratios (SNRs), as demonstrated in Supplementary Fig. S5, and stabilized once sufficient target observations were included (Supplementary Fig. S6). Slight performance degradation was observed for models near the edges of the FOV, likely reflecting lower SNR and stronger PSF distortions. Taken together, these results indicate that DEPR recovers the underlying ePSF with high fidelity throughout the imaging volume and maintains reliable performance under practical conditions.

The ePSF recovered by DEPR provides a transferable system model that can be applied to deconvolution and cross correlation, and is especially useful for localization. By fitting the target intensity profiles against the learned ePSF, it yields more accurate sub-pixel centroids and thereby improves overall super-resolution image quality. We evaluated localization accuracy gains by generating sub-pixel point targets along the x, y, and z axes, and localizing them with both CG and DEPR-based LSF, which shows reduced systematic bias and tightened residuals, with the largest benefit in the axial direction where PSF asymmetry is most pronounced (Fig. 3f). Simulated microtubule reconstructions (see methods for details) further showed better preservation of filament continuity, finer details, and sharper profiles than conventional CG-based approach (Fig. 3g and 3h). Localization histograms also showed markedly better agreement with the simulated reference distribution when using DEPR, indicating a reduction in position dependent bias (Fig. 3i). Overall, these results demonstrate that DEPR enables accurate *in-situ* ePSF estimation under realistic conditions, which further translates into localization enhancement.

### Large-scale *in vivo* LOT microangiography

Building on DEPR, a modified *in vivo* localization strategy was developed and validated. Conventional approaches use morphological thresholding (MT) and centroid estimation, henceforth referred as MT-CG, which removes small targets to improve image quality but reduces data utilization^20,47^. The proposed alternative uses regional-maxima (RM) detection without morphological constraints, preserving more candidates but requiring higher sub-pixel accuracy. This is achieved by fitting intensity profiles to the retrieved field-dependent ePSFs, yielding the RM-LSF strategy. Supplementary Fig. S7 depicts the MT-CG and RM-LSF workflows. Dynamic volumetric LOT of mouse brain vasculature in both NIR-I and NIR-II windows enabled direct comparison of the two strategies.

In the first experiment, dichloromethane (DCM) microdroplets were intravenously injected and imaged at 780 nm excitation. Their sub-capillary size enabled free circulation without vascular arrest. Fig. 4a shows volumetric vascular maps reconstructed from particle localizations using conventional MT-CG and DEPR-based RM-LSF strategies, where color encoding indicates depth and clearly separates cerebral from calvarial structures. A representative *in vivo* ePSF from the central FOV is shown in Fig. 4b, with orthogonal cross-sections and axis profiles confirming accurate system characterization. Spatial variation of ePSFs across the FOV further validated the field-dependent modeling (Supplementary Fig. S8). Zoomed projections (Fig. 4c) and differential images (Fig. 4d) demonstrate more and better-defined vascular details under identical frame counts by the proposed method. Object counts and line profiles (Fig. 4e and 4f) show that by eliminating the MT step, it achieves over 50% higher data utilization and produces narrower features with desirable contrast.

**Figure 4.**
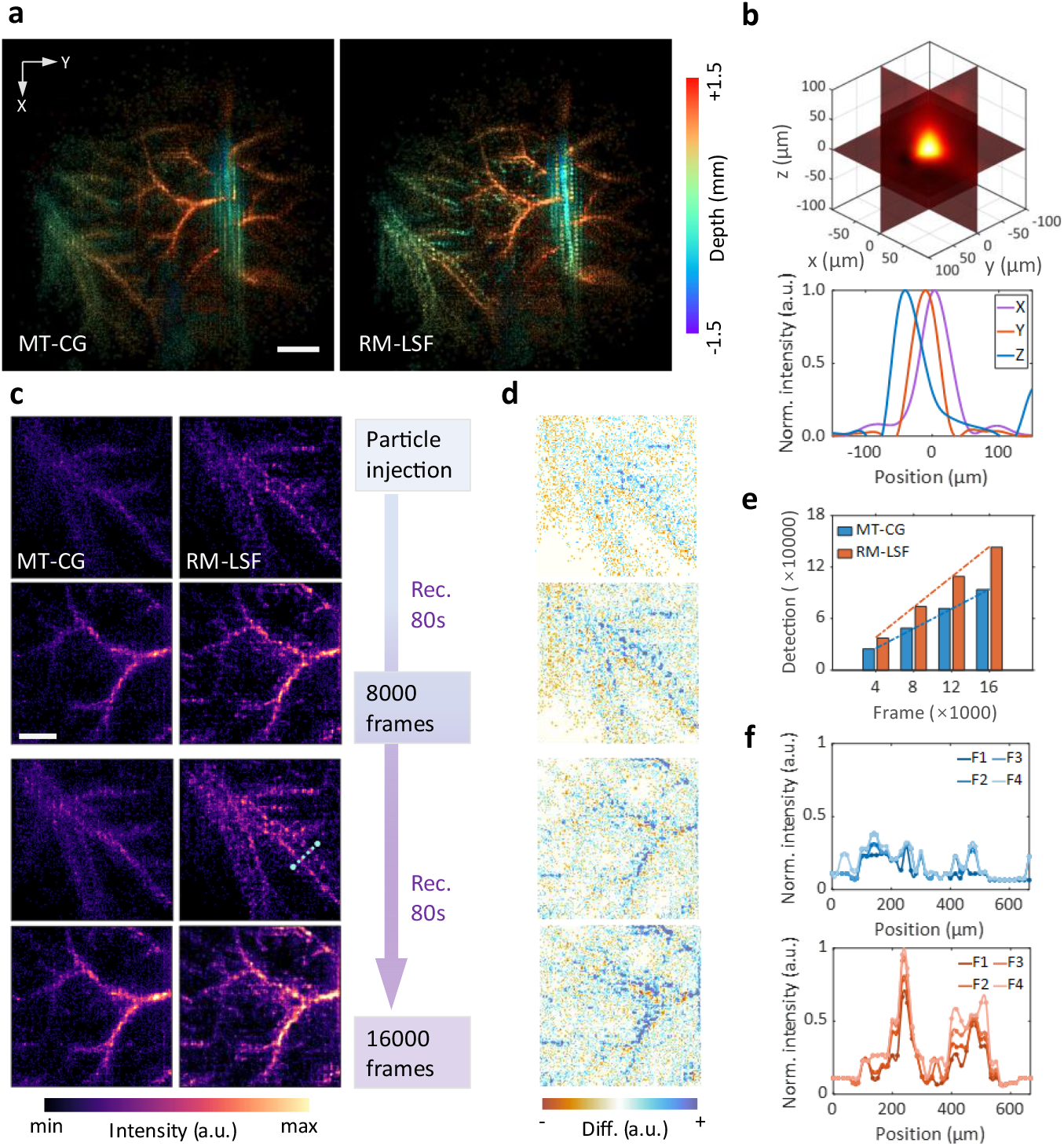
*In vivo* LOT imaging of mouse brain vasculature under NIR-I excitation with DCM microparticles. **a**, Volumetric vascular maps rendered from particle localizations using conventional morphological thresholding with center-of-gravity (MT-CG, left) and DEPR-based regional-maxima detection with least-squares fitting (RM-LSF, right) strategies. Color encoding represents depth information revealing cerebral and calvaria structures. Scalebar – 1 mm. **b**, Representative *in vivo* ePSF retrieved from the central FOV region showing volumetric orthogonal cross-sections (top) and axis profiles along x, y and z directions (bottom). **c**, Zoomed regional MIPs comparing conventional MT-CG (left column) and DEPR-based RM-LSF (right column) strategies at 8000 and 16000 frames. Scalebar – 500 μm, a.u. – arbitrary units. **d**, Differential images highlighting enhanced vascular details achieved by the DEPR-based method. **e**, Quantitative analysis showing number of identified objects under same number of recorded frames demonstrating higher data-utilization efficiency. **f**, Line profiles across vascular features comparing conventional (blue) and DEPR-based (red) methods at different frame counts, demonstrating enhanced contrast and spatial resolution. F1 – 4000 frames, F2 – 8000 frames, F3 – 4000 frames, F4 – 8000 frames.

To demonstrate broader applicability, additional LOT imaging was performed with CaCO_3_@CuS/PDA microparticles absorbing light at 1064 nm in the NIR-II window. These absorbers generated strong and stable OA signals, enabling more vascular detail with fewer frames compared to DCM droplets^48^. Fig. 5a presents superimposed volumetic reconstructions, where finer structures are consistently resolved by the proposed method for different frame amounts (Supplementary Fig. S9). A representative *in vivo* ePSF is shown in Fig. 5b, distinct from the previous case, reflecting altered optical and acoustic properties. Enlarged regions (Fig. 5c and 5d) and line profiles (Fig. 5e) confirm sharper vessel depiction and clear peak separation. FWHM analysis demonstrated significant reductions, from 212.3 µm to 84.6 µm and from 75.9 µm to 41.5 µm, highlighting improved resolution. Beyond structural imaging, the enhanced sub-pixel precision of the DEPR-based RM-LSF strategy provides a robust basis for functional analysis. As an example, particle tracking was performed using localized absorbers to generate a vascular flow map (Fig. 5f)^49^, which revealed spatially resolved trajectories across the cerebral network. This proof-of-concept highlights the potential of the method to extend beyond static reconstructions toward dynamic functional readouts such as flow quantification and perfusion analysis.

**Figure 5.**
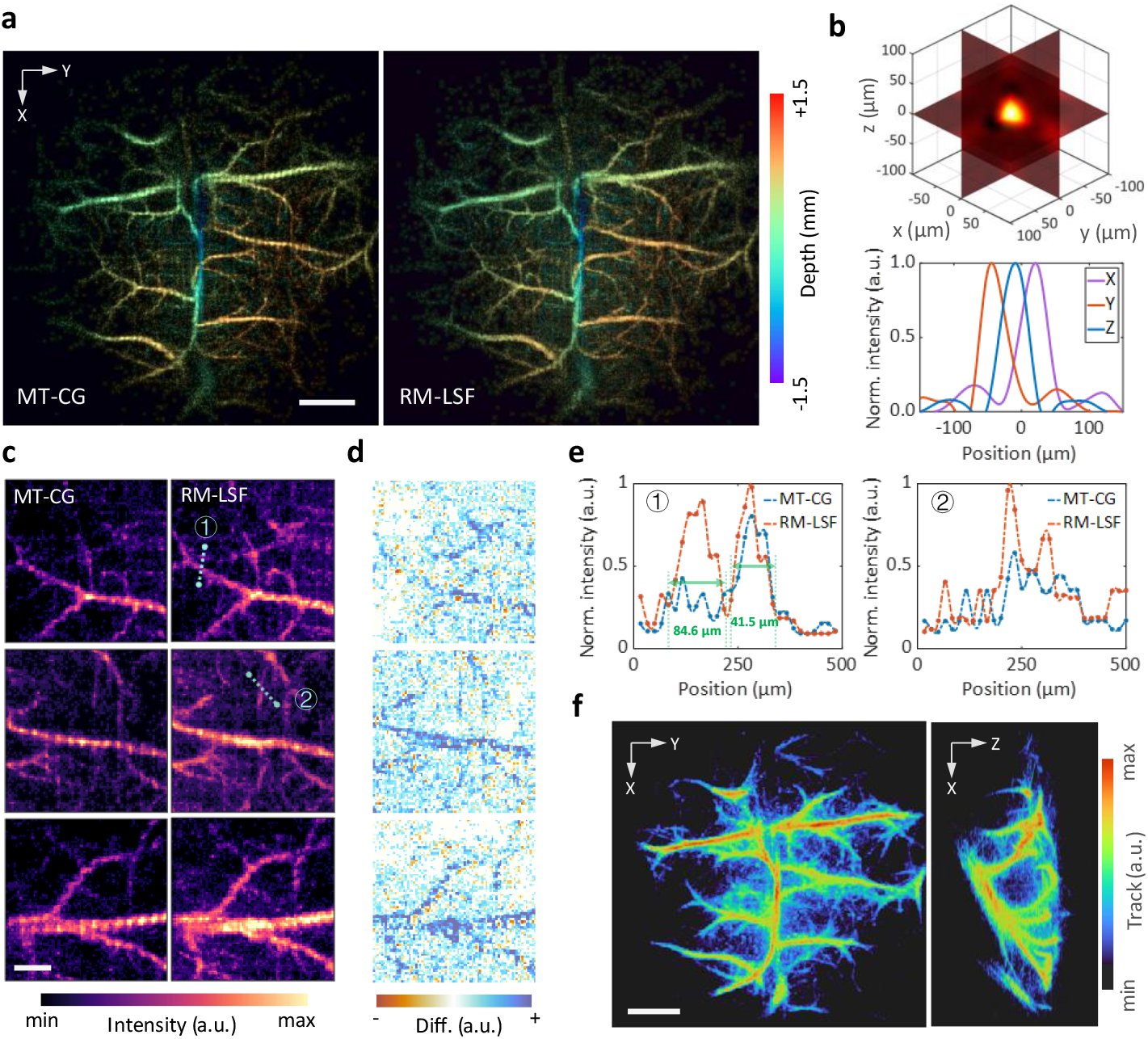
*In vivo* LOT imaging of mouse brain vasculature under NIR-II excitation with CaCO_3_@CuS/PDA microparticles. **a**, Volumetric vascular maps rendered from particle localizations using conventional MT-CG (left) and DEPR-based RM-LSF (right) strategies under 1064 nm excitation. Color encoding represents depth information. Scalebar – 1 mm. **b**, Representative *in vivo* ePSF retrieved from the central FOV region showing volumetric orthogonal cross-sections (top) and axis profiles along x, y and z directions (bottom), displaying distinct morphology from NIR-I imaging. **c**, Zoomed regional comparisons showing enhanced vascular details with conventional (left column) and DEPR-based (right column) methods under same frame counts. Scalebar – 300 μm, a.u. – arbitrary units. **d**, Differential images highlighting improved microvascular details achieved by DEPR-based method. **e**, Line profiles across vascular structures demonstrating peak separation capabilities, with FWHM reductions from 212.3 µm to 84.6 µm, and from 75.9 µm to 41.5 µm for selected vessels. **f**, Particle tracking analysis based on the DEPR-based RM-LSF localization output, generating tracking maps of cerebral vasculature (left: axial view, right: sagittal view) demonstrating the method’s potential for dynamic functional imaging applications. Scalebar – 1 mm.

To further evaluate the impact of localization accuracy on image quality, another baseline strategy, referred as RM-CG, was implemented, where absorbers identified by RM detection were directly localized using centroid estimation without the MT procedure. LOT images reconstructed with RM-CG displayed more pronounced gridded artifacts compared to the DEPR-based RM-LSF approach, as shown in the enlarged vascular regions and differential images in Fig. 6a. In contrast, RM-LSF consistently provided sharper vascular structures and substantially reduced artifacts for both NIR-I and NIR-II windows. Two-dimensional Fourier spectra of reconstructed images (Fig. 6b) showed a clear reduction of spectral peaks corresponding to periodic grid artifacts in the final reconstructions. Sub-pixel localization histograms from the NIR-I dataset (Fig. 6c) further demonstrated that RM-LSF yielded a more uniform positional distribution than RM-CG, mitigating clustering at specific sub-pixel sites. Furthermore, the proposed strategy introduced only a modest computational overhead, while total runtime was still dominated by image reconstruction (Supplementary Fig. S10). For the NIR-II dataset, radial power spectral density (PSD) analysis (Fig. 6d) confirmed attenuation of regularly distributed spectral components. Quantitative frequency-domain analysis showed that RM-LSF suppressed mid-to-low frequency ratios and spectral centroids by 25.8% and 20.1%, respectively, indicating superior noise rejection. Together, these results establish DEPR-based RM-LSF as a reliable and high-fidelity localization strategy for *in vivo* vascular imaging.

**Figure 6.**
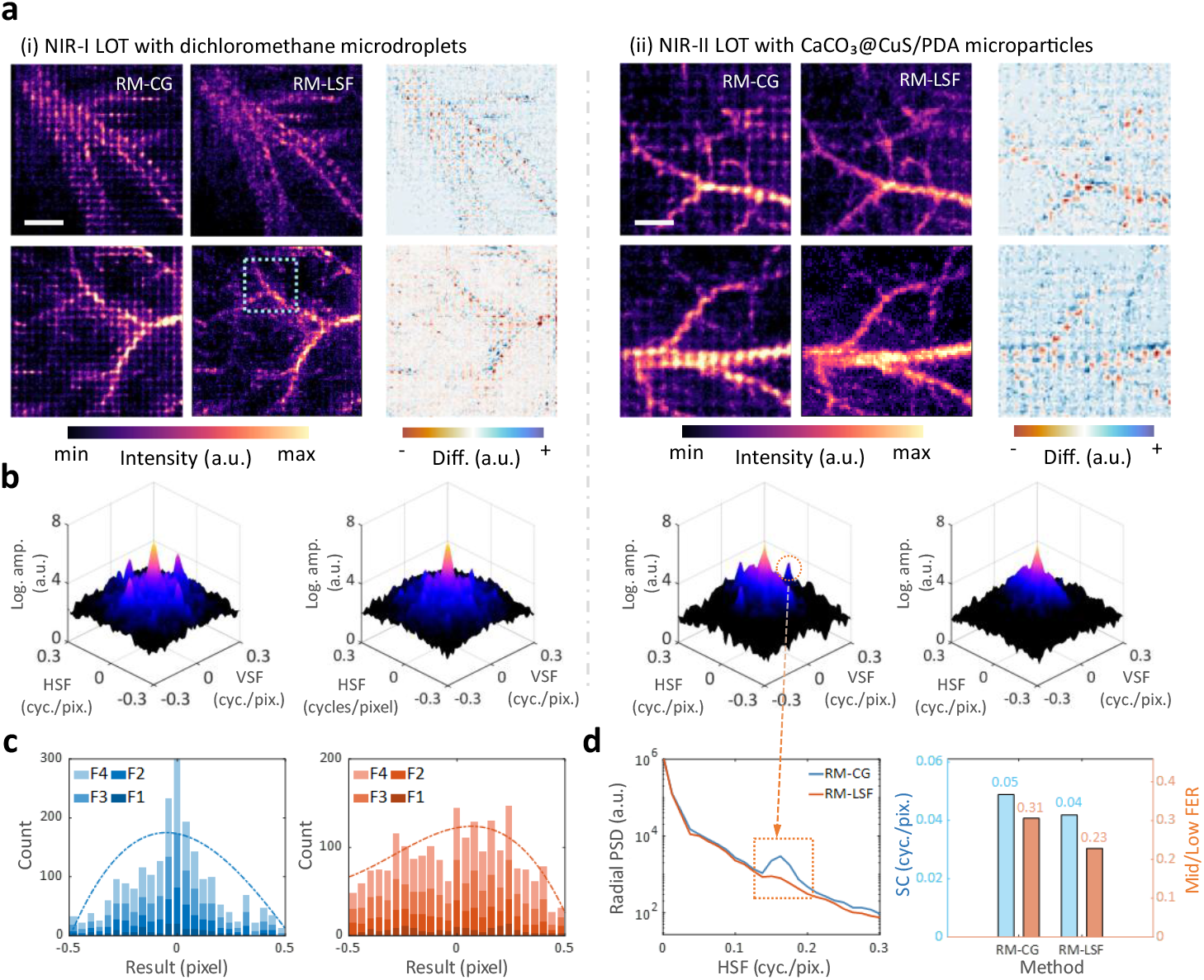
Comparative analysis of regional-maxima detection with direct center-of-gravity (RM-CG) estimation and the proposed RM-LSF strategies *in vivo*. **a**, Super-resolved LOT images from NIR-I and NIR-II datasets reconstructed with RM-CG and RM-LSF, together with differential maps. RM-LSF consistently sharpens vascular details and suppresses grid artefacts compared to RM-CG. (i) Scalebar – 500 μm, (ii) Scalebar – 300 μm, a.u. – arbitrary units. **b**, Two-dimensional Fourier spectra of reconstructed images. RM-LSF effectively reduces the spectral peaks corresponding to periodic grid artifacts in the final reconstructions. HSF – horizontal spatial frequency, VSF – vertical spatial frequency, cyc./pix. – cycles per pixel. **c**, Sub-pixel localization histograms within the representative square region from the NIR-I dataset showing that RM-LSF yields more uniform distributions, mitigating clustering effects seen with RM-CG. F1 – 4000 frames, F2 – 8000 frames, F3 – 4000 frames, F4 – 8000 frames. **d**, Radial power spectral density (PSD) analysis and quantitative metrics from the NIR-II dataset confirm superior noise rejection, as the RM-LSF method suppressed mid-to-low frequency ratios and spectral centroids by 25.8% and 20.1%, respectively. SC – spectral centroid, FER – frequency energy ratio.

## Discussion

This study introduces a data-driven framework that retrieves ePSFs directly from experimental data, enabling optimized localization under realistic imaging conditions. By reconstructing continuous models from discretized measurements, the method transforms pixelated responses into smooth ePSFs that preserve sub-pixel features and reduce systematic bias. This allows precise localization in under-sampled conditions, thus overcoming the challenging trade-off between resolution and FOV while reducing computational demands. Statistical robustness from aggregating thousands of localizations is particularly effective in low-SNR environments. The *in-situ* modeling captures field-dependent variations and sample-specific aberrations, yielding more faithful vascular reconstructions. Benefiting from DEPR-based enhancement, the method mitigates artifacts that commonly affect LOT. Moreover, by eliminating the MT procedure, it substantially increases data utilization efficiency, allowing more absorbers to contribute to the reconstruction and producing denser, higher-fidelity vascular maps from fewer acquired frames.

The method’s performance relies on adequate spatial distribution and depth range of point sources, as insufficient sampling diversity can lead to incomplete PSF models, particularly in the axial dimension. Reconstruction fidelity may also be affected at large depths due to cumulative aberrations, or when absorbers are positioned too closely to be distinguished. This could be addressed through delicate transducer design or specific algorithmic refinements^50,51^. Herein, we employed LSF for absorber localization, which offers computational simplicity and efficiency, while future implementations may integrate maximum-likelihood estimation (MLE) to further push the boundaries for optimal precision^52^.

The data-driven philosophy underlying DEPR naturally aligns with modern machine learning paradigms. It operates in a self-supervised manner, as localization labels are not externally provided but instead estimated and iteratively refined within the framework. While the current implementation reconstructs ePSFs from explicitly defined statistical features, deep learning architectures could directly learn optimal representations from raw data, capturing more complex relationships between measurements and absorber positions. Physics-informed neural networks are particularly promising, as they combine the interpretability of model-based approaches with the adaptability of learning-based methods to ensure physically meaningful representations^53^. Such extensions could address current limitations in localization performance and enable real-time processing through GPU acceleration, paving the way for broader applicability in advanced imaging.

Demonstrated here for OA imaging, the DEPR framework generally addresses fundamental challenges shared by all localization-based super-resolution imaging modalities. Issues such as PSF distortion, spatial sampling discretization, and position-dependent bias are universal across imaging systems, including fluorescence microscopy, ultrasound localization microscopy, and astronomical imaging. By retrieving system-specific responses directly from experimental data without prior assumptions, DEPR offers a broadly applicable strategy for imaging through complex media where theoretical models fall short. Its generalizability positions it as a powerful tool for improving imaging performance across diverse platforms, with potential to drive breakthroughs in biomedical research, clinical diagnostics, and large-scale observational sciences.

## Methods

### OA tomography system

OA tomographic imaging of the mouse brain was performed using a custom spherical array system (Fig. 1a) comprising 512 ultrasound transducers arranged hemispherically at 40 mm radius with 150° angular coverage (Imasonic SaS, France)^20^. The trapezoidal elements (∼3.3×3.8 mm^2^), each operating at 7 MHz center frequency with >80% detection bandwidth, were uniformly distributed across 13 concentric rings. The array design incorporated a central 8 mm cylindrical aperture and three 4 mm lateral apertures positioned at 45° elevation and separated by 120° azimuthally.

Pulsed laser excitation (<10 ns) was delivered at wavelengths optimized for NIR-I and NIR-II windows, matched to microparticle absorption peaks. A custom four-arm fiber bundle (CeramOptec GmbH) routed through the array apertures provided uniform cortical illumination at ∼20 mJ/cm^2^ fluence. OA signals underwent 40 dB amplification before digitization at 40 MS/s (Falkenstein Mikrosysteme GmbH) and transfer for reconstruction.

Signal preprocessing comprised bandpass filtering (1–7 MHz) and SVD-based clutter suppression. Volumetric images covering the specified region were reconstructed with 0.1 mm isotropic pixels (e.g., 60×60×60 volumetric pixels spanning 6×6×6 mm^3^) using GPU-accelerated standard BP algorithm. This volumetric pixel size intentionally operated in the sub-Nyquist regime to balance computational efficiency with spatial coverage. High-frequency components were preserved to maximize native resolution, while temporal tracking of microparticle dynamics enabled super-resolution LOT reconstruction.

### DEPR implementation pipeline

DEPR is formulated as an inverse problem in which discrete ROI measurements from transient point sources inform a continuous and field-dependent ePSF that best explains observations across sub-pixel positions. The framework treats each detected point source response as a feature patch and extracts stable features by pooling many such patches across time and space, thereby learning the underlying system response directly from data without external calibration. Let *I* _*k*_ ∈ *R*^*N* ×*N* ×*N*^ denote the volumetic ROI around the k-th point source arising from an unknown source position. The source position decomposes into an integer volumetric pixel center 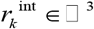 and a sub-pixel offset *δ*_*k*_ ∈[−0.5,0.5), *k* ∈ *x, y, z*, such that the true position equals

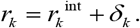

The continuous ePSF *h* (*r*) relates to observed intensities through

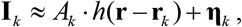

Where *A*_*k*_ is the point source amplitude, and *η*_*k*_ denotes additive noise. The objective is to estimate *h* (*r*) together with improved point source positions using many observations {*I*_*k*_ } sampled at diverse *δ*_*k*_ across the imaging volume.

To account for field dependence, the imaging volume is partitioned into subregions {*R*_*m*_}, within which the ePSF is assumed locally stationary. Each localized emission event *I*_*k*_ is assigned to its corresponding subregion *R*_*m*_ according to its location 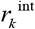.

ROIs are then re-centered within a fixed window Ω ⊂ □^3^ around 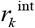, which also plays the role of feature extraction by isolating the local intensity pattern associated with a single point source. Sub-pixel offsets *δ*_*k*_ are typically estimated using the CG method, and GF is applied selectively to enhance robustness under moderate SNR. The CG estimator is defined as

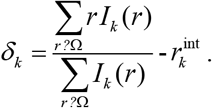

The estimated offsets are quantized into bin indices *b*_*k*_ with a chosen bin resolution Δ∈(0, 0.5 ], so that

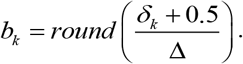

All ROIs that fall into the same sub-pixel bin are grouped for statistical aggregation. Let *I*_*b*_ (*r*) be the set of intensity values at pixel position *r* from all ROIs sharing bin *b*. To mitigate the effect of outliers, robust averaging is applied with statistical filtering as

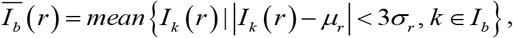

where *μ*_**r**_, *σ*_**r**_ are the sample mean and standard deviation at pixel *r*, respectively. The result is a collection of the averaged ROIs 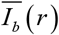 that capture sub-pixel sampled responses.

Continuous ePSFs *h*_*m*_ (*r*) for each subregion *R*_*m*_ are reconstructed by cubic spline interpolation applied to the aggregated measurements

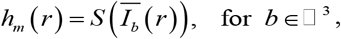

where *S* denotes the spline operator. This interpolation recovers genuine sub-pixel structure from pixelated measurements, transforming discrete observations into smooth, continuous representations that preserve fine spatial details lost in conventional approaches.

The framework implements self-supervised refinement through iterative optimization. Reconstructed ePSFs enable improved localization via LSF

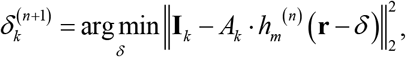

where 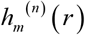 is the reconstructed ePSF at iteration ***n***. The updated positions 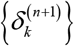 are fed back into the binning process, progressively refining both the ePSF models and localization accuracy. This self-supervised strategy requires no external labels, instead leveraging the internal consistency of the data to converge toward optimal representations, typically within 3–4 cycles.

The final output is a library {*h*_*m*_ (*r*)} of high-resolution ePSFs indexed by subregion *R*_*m*_. The central FOV typically yields the most statistically reliable model due to highest point source density, serving as a default template *h*_0_ (*r*) for regions with insufficient sampling or as an initialization prior for subsequent datasets. By combining feature identification and aggregation with iterative self-supervised updates, DEPR bridges under-sampling and accurate sub-pixel localization for realistic *in situ* conditions.

### Analysis of calibration and simulation data

To quantify system response variations across the field, PCA-based similarity metric was implemented to the calibrated data^46^. Each ePSF was vectorized and projected onto the first nine principal components, which together captured more than 95% of the variance. Spatial deviation was quantified by calculating the Euclidean distance between the projection coefficients of each ePSF and those of the central reference. These distances were mapped to their corresponding field positions, revealing smooth model evolution across the imaging volume (Fig. 3c).

To benchmark reconstruction fidelity, synthetic datasets were generated using experimentally measured ground truth PSFs. Approximately 60000 targets were placed at random sub-pixel positions, each rendered as a low-resolution pixelized observation by sampling its corresponding PSF with a prescribed offset. Multiplicative noise (2%) was added to mimic realistic acquisition conditions. The resulting dataset, consisting of thousands of noisy, discretized ROIs, was processed using the DEPR pipeline, including sub-pixel binning, statistical filtering, spline interpolation, and iterative least-squares refinement. Reconstruction accuracy was evaluated by comparing recovered ePSFs against ground truth reference, and residuals were quantified at the pixel level after normalization.

To evaluate localization performance, synthetic point targets were generated with known sub-pixel positions uniformly spanning –0.5 to +0.5 volumetric pixels along the x, y, and z axes (101 positions per axis, 50 replicates each). Localization was performed using both the conventional CG method and LSF with DEPR-derived ePSFs. Localization error was defined as the absolute difference between estimated and ground truth positions. Residuals were summarized as mean ± standard deviation and plotted against sub-pixel position.

To demonstrate structural reconstruction enhancement, volumetric microtubule networks were simulated with approximately 60000 targets distributed along tubular trajectories. Each was convolved with a high-resolution ground truth PSF (501×501 ×501 volumetric pixels) at its specified sub-pixel position, and under-sampled images (5×5×5 volumetric pixels) were generated with 2% multiplicative noise. Synthetic datasets were then processed with both the CG-based and DEPR-based pipelines. Super-resolved volumes were reconstructed by mapping localized positions into an up-sampled grid (factor 20) and applying Gaussian filtering (σ=0.6). Lateral intensity profiles and reconstructed filament continuity were analyzed to compare contrast and resolution between methods.

### *In vivo* experiments

As NIR-I absorbers, ∼5 µm dichloromethane (DCM) microdroplets were used. These contained IR-780 dye at approximately 200 mM concentration with an extinction coefficient of ∼250000 M^−1^cm^−1^, which corresponds to a per-particle absorption three to four orders of magnitude higher than red blood cells^20^. The experiment employed a short-pulsed laser at 780 nm wavelength and a spherical array of 512 ultrasound transducers for imaging at a frame rate of 100 Hz (100 volumetric images per second). A nude-Fox1nu mice was administered with 100 µL of microdroplet emulsion via tail vein injection, with a total acquisition time of 420 seconds.

As NIR-II absorbers, CaCO_3_@CuS/PDA microparticles with ∼3.6 μm average diameter were used for LOT imaging^48^. These particles were created by loading copper sulfide nanoparticles (9-10nm) onto calcium carbonate microparticles and coating them with polydopamine. The experiment utilized a 1064 nm wavelength laser (NIR-II window) and a similar spherical ultrasound array at 100 Hz frame rate, with a total acquisition time of 360 seconds (36000 frames). A 5-8 week-old nude mice was administered with 100 μL of microparticle suspension (2×10^8^ particles/mL) via tail vein injection. These particles could form images of major vessels in a relatively short time (about 30 seconds), with complete microvascular networks visible after 330 seconds.

All animal experiments were performed in accordance with the Swiss Federal Act on Animal Protection and were approved by the Cantonal Veterinary Office of Zurich. Mice were maintained in individually ventilated, temperature-controlled cages under a 12-hour light/dark cycle, with *ad-libitum* access to pelleted food (3437PXL15, Cargill) and water. For imaging procedures, animals were anesthetized with isoflurane (5% v/v for induction and 1.5% v/v for maintenance; Abbott, Cham, Switzerland) delivered in an oxygen/air mixture (100/400 mL/min).

## Supplementary Information

Supplementary Information is available from the attachment file.

## Data Availability Statement

The data that support the findings of this study are available from the corresponding author upon reasonable request.

## Acknowledgements

This work was supported by the National Nature Science Foundation of China (No. U24A6006), the National Key Research and Development Program of China (No. 2023YFB3906300), the Innosuisse – Swiss Innovation Agency (grant 51767.1 IP-LS), and the Swiss National Science Foundation (grant 310030_192757).

