## Supplementary Information for "Data-Driven Retrieval of Effective Point Spread Functions for Super-Resolution Optoacoustic Imaging"

**This file includes:**

**Note S1.** Principles of optoacoustic tomography back-projection reconstruction and sources of bias in center-of-gravity localization.
**Note S2.** Analysis of optimal sub-pixel binning interval for DEPR.
**Note S3.** Impact of iteration rounds on ePSF reconstruction accuracy.
**Note S4.** Time-frequency analysis of the field-dependent system response.
**Note S5.** Validation of DEPR robustness against varying noise conditions.
**Note S6.** Impact of the number of targets on reconstruction fidelity.
**Note S7.** Schematic diagram of the baseline and the modified localization workflow.
**Note S8.** *In vivo* characterization of field-dependent ePSFs.
**Note S9.** Comparison of the rendered LOT image with respect to cumulative frames.
**Note S10.** Computational performance comparison of localization methods.

**Note S1. Principles of optoacoustic tomography back-projection reconstruction and sources of bias in center-of-gravity localization.**

The system layout for localization optoacoustic tomography (LOT) is presented in Fig. S1, which is consistent with conventional optoacoustic tomography (OAT) architectures and offers broad compatibility with any system supporting real-time imaging.

The fundamental principle of back-projection reconstruction is to determine the initial energy absorption field $H(r_0)$ from the pressure waves $p(r, t)$ generated within a medium following pulsed laser illumination. This process is governed by the optoacoustic wave equation[1]:

$$\frac{\partial^2 p(r,t)}{\partial t^2} - c^2 \Delta p(r,t) = \Gamma H(r,t) \frac{\partial \delta(t)}{\partial t} \tag{1}$$

where $c$ represents the speed of sound, $\Gamma$ is the Gruneisen parameter, and $H(r, t)$ is the absorbed energy density. For excitation pulses of sufficiently short duration, $H(r, t)$ can be approximated as $H(r_0)\delta(t)$. The forward solution, derived from the Poisson solution for the wave equation, describes the pressure field as:

$$p(r,t) = \frac{\Gamma}{4\pi c} \frac{\partial}{\partial t} \int_{S_0} \frac{H(r_0)}{|r - r_0|} dS_0 \tag{2}$$

The goal of optoacoustic reconstruction is the inverse problem of determining $H(r_0)$ from the recorded $p(r, t)$. This is commonly achieved using the universal back-projection algorithm, which in its continuous form is approximated as[2]:

$$H(r_0) = \frac{1}{\Gamma} \int_{\Omega} \frac{d\Omega}{\Omega} \left[ 2p(r,t) - 2t \frac{\partial p(r,t)}{\partial t} \right] \Big|_{t=\frac{|r-r_0|}{c}} \tag{3}$$

In practical computation, this formula is numerically discretized. For a specific transducer position $r_i$ and a reconstruction volumetric pixel $r_0^j$, the discretized formula is[3]:

$$H(r_0^j) = \sum_i \left[ p(r_i, t_{ji}) - t_{ji} \frac{\partial p(r_i, t_{ji})}{\partial t} \right] \tag{4}$$

$$t_{ji} = \frac{\left| r_i - r_0^j \right|}{c} \tag{5}$$

where $t_{ji}$ is the time-of-flight from $r_i$ to $r_0^j$.

This algorithm is often simplified for computational efficiency by neglecting the first term, which is comparatively small, and omitting the multiplication by time, which varies negligibly within the region of interest. The simplified and practical reconstruction formula thus becomes:

$$H(r_0^j) = \sum_i \left[ -\frac{\partial p(r_i, t_{ji})}{\partial t} \right] = \sum_i p_{filt}(r_i, t_{ji}) \tag{6}$$

where $p_{filt}(r_i,t_{ji})$ represents the signal after appropriate filtering and differentiation.

In LOT, the precise target positions are estimated subsequent to image reconstruction. The center-of-gravity (CG) method is a widely-adopted approach valued for its simplicity and speed[4]. It calculates the centroid coordinates by operating on the discrete volumetric pixel values $P_{ijk}$ within a defined region of interest (ROI) using the summations:

$$(x_{CG},y_{CG},z_{CG})=(\frac{\sum_{ROI}iP_{ijk}}{\sum_{ROI}P_{ijk}},\frac{\sum_{ROI}jP_{ijk}}{\sum_{ROI}P_{ijk}},\frac{\sum_{ROI}kP_{ijk}}{\sum_{ROI}P_{ijk}}) \tag{7}$$

However, this method harbors an intrinsic inaccuracy. This error arises from the fundamental discrepancy between the discrete calculation performed on sampled data and the continuous nature of the underlying instrumental point spread function (PSF) *I(x,y,z)*. The true centroid is properly defined by the continuous integrals:

$$(x^*,y^*,z^*)=(\frac{\iiint_{ROI}xI(x,y,z)dxdydz}{\iiint_{ROI}I(x,y,z)dxdydz},\frac{\iiint_{ROI}yI(x,y,z)dxdydz}{\iiint_{ROI}I(x,y,z)dxdydz},\frac{\iiint_{ROI}zI(x,y,z)dxdydz}{\iiint_{ROI}I(x,y,z)dxdydz}) \tag{8}$$

Therefore, the localization error can be exemplified by:

$$(e_x,e_y,e_z)=(x_{CG},y_{CG},z_{CG})-(x^*,y^*,z^*) \tag{9}$$

which stems directly from the fact that the discrete summation over pixels is merely an approximation of the continuous integration over the true signal distribution. Notably, as discussed in our previous studies[5], the impact of this deviation becomes increasingly significant as the PSF size decreases, thereby substantially affecting the sub-pixel localization accuracy.

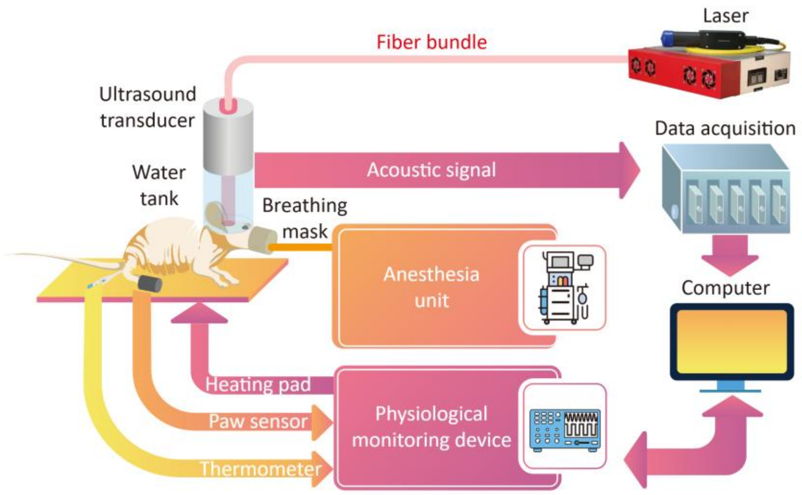


**Figure S1. Experimental setup for *in vivo* optoacoustic tomography (OAT).**
Schematic of the OAT system used for acquiring localization optoacoustic tomography (LOT) images of the murine brain, detailing the laser illumination, acoustic signal detection, and physiological monitoring components.

**Note S2. Analysis of optimal sub-pixel binning interval for DEPR.**

To investigate the optimal sub-pixel binning interval ($\Delta$) for the data-driven effective point spread function retrieval (DEPR) framework, simulation study was performed to evaluate reconstruction accuracy with respect to this parameter. The choice of $\Delta$ directly impacts the fidelity of the reconstructed effective point spread function (ePSF) by balancing sampling resolution against statistical robustness. Four distinct intervals were tested: 1/3, 1/4, 1/5, and 1/6 pixels, with the results shown in Fig. S2.

Clear trade-off was observed related to the interval selection. Finer sampling intervals, such as $\Delta$=1/5 or 1/6 pixels, theoretically offer higher sub-pixel resolution. This allows the ePSF model to capture finer features and produce profiles closer to the ground truth by reducing the energy-averaging effect within each bin. However, this sensitivity comes at a cost. Finer intervals are more susceptible to the influence of initial localization errors, where target incorrectly assigned to an adjacent bin during the iterative process can introduce artifacts into the model, as evidenced by the increased peripheral error in the $\Delta$=1/6 pixels interval reconstruction. Conversely, a coarser interval, such as 1/3 pixels, provides greater statistical robustness by averaging more target observations per bin. This, however, reduces the effective sub-pixel resolution, and the resulting ePSF relies more heavily on interpolation than on aggregated data, yielding an overly smooth model that fails to capture the true system response (Fig. S2a and b).

This trade-off was further evaluated by applying the resulting ePSF models to the reconstruction of a synthetic microtube network (Fig. S2c). The intensity profiles demonstrate each model's capability to resolve closely spaced structures (Fig. S2d). The results derived from the $\Delta$=1/4 pixels interval yielded the best overall performance, demonstrating sharp, well-defined peaks and better resolving capability between adjacent filaments, which suggests it provides the best imaging performance by balancing the competing factors. We therefore selected 1/4 pixels as the optimal interval for our study, as it achieves the best reconstruction effect by maximizing fidelity while maintaining robustness against initial localization errors.

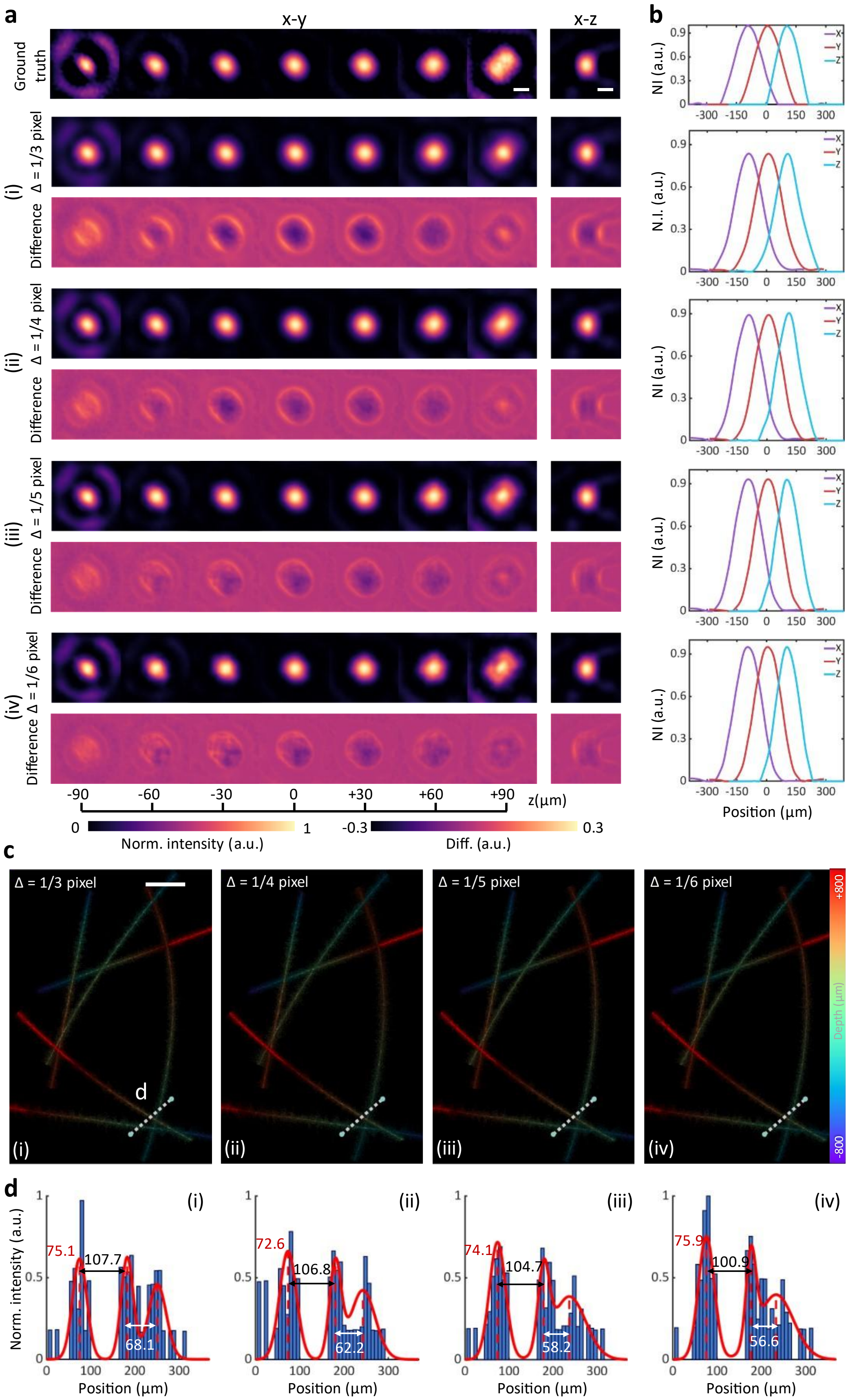


**Figure S2. Optimization of the sub-pixel binning interval for data-driven effective point**

**spread function retrieval (DEPR).**

**(a)** Axial (x-y) and sagittal (x-z) views of the ground truth and the effective point spread functions (ePSFs) reconstructed using different sampling intervals (Δ = 1/3, 1/4, 1/5, and 1/6 pixels). Difference maps are shown below each corresponding model to visualize the distribution of model bias. Scalebar – 100 µm, NI – normalized intensity, a.u. – arbitrary units. **(b)** Normalized intensity profiles along the x, y, and z axes for the corresponding reconstructed ePSFs shown in (a). For clearer comparison, the profiles are spatially offset by 100 µm to prevent overlap between the axes. **(c)** Reconstructions of a synthetic microtube network using the ePSF models derived from the different sampling intervals in (a). Scalebar – 200 µm. **(d)** Normalized intensity profiles (blue histograms) and fitted profiles (red curves) along the dashed white lines indicated in (c). The relative positions (red and white labels) and spacings (black labels) of the fitted peaks reflect the reconstruction accuracy and resolving capability of each model.

**Note S3. Impact of iteration rounds on ePSF reconstruction accuracy.**

To evaluate the convergence behavior of the self-supervised DEPR framework, the relationship between ePSF reconstruction accuracy and the number of refinement iterations was analyzed. Using the simulation dataset and the optimized 1/4 pixels binning interval, the ePSF model and the corresponding error distribution over four successive iterations was established.

As shown in Fig. S3a, the reconstruction accuracy of the ePSF model improves with each iteration. More substantial refinement occurs from the first to third iteration rounds, where both the central core and the peripheral structure of the ePSF become more clearly defined, and the corresponding intensity difference is visibly reduced. The model's change in accuracy begins to stabilize afterwards, with negligible improvements observed after the fourth iteration.

This convergence is quantitatively confirmed by the model's residual distribution histograms in Fig. S3b. The histograms, which represent the pixel-wise normalized intensity errors, become progressively narrower and more sharply peaked around zero, indicating a reduction in overall model error. The minimal change between iteration 3 and 4 confirms that the refinement process has effectively converged. This simulation demonstrates that the self-supervised DEPR method possesses reliable convergence characteristics. For practical applications, an appropriate number of iterations should be selected based on the experimental setup and imaging region to balance model fidelity and computational cost.

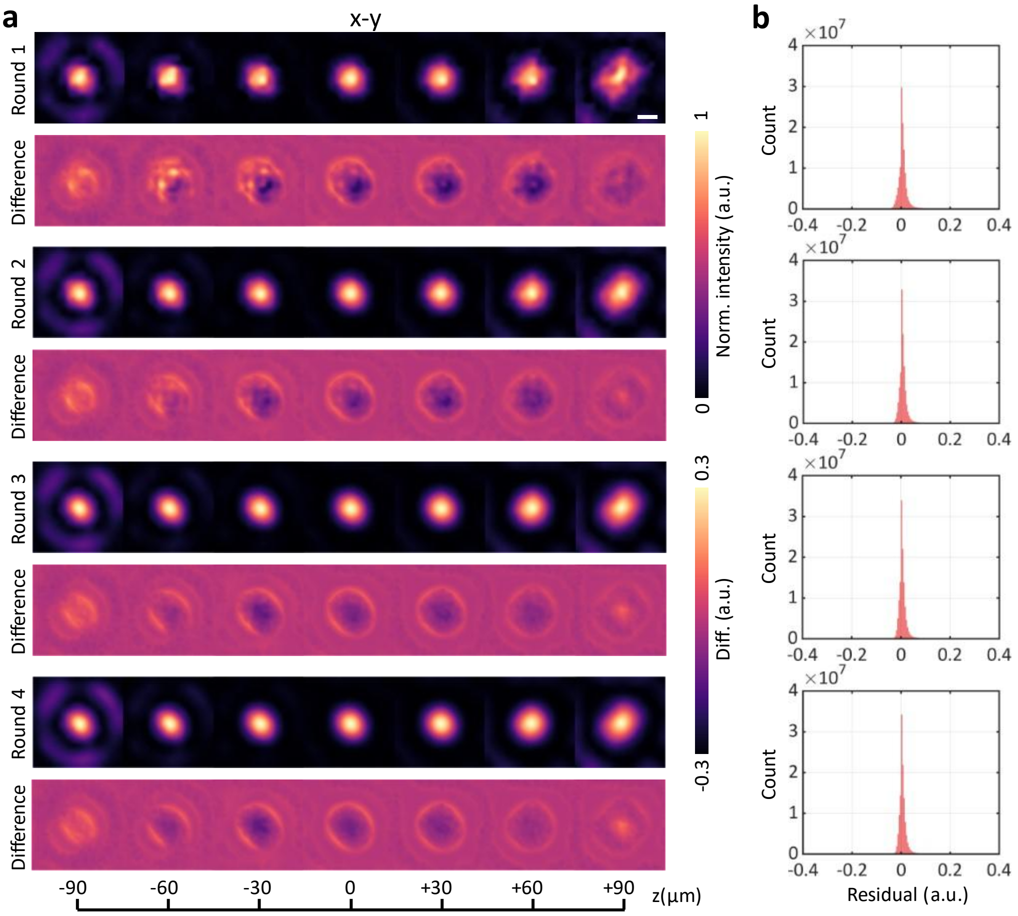


**Figure S3. Convergence behavior of the iterative DEPR framework.**

**(a)** Axial (x-y) views of the reconstructed ePSF models shown after 1, 2, 3, and 4 iteration rounds. Corresponding difference maps are displayed below each model to visualize the reduction in intensity error. Scalebar – 100 µm, a.u. – arbitrary units. **(b)** Histograms of the pixel-wise normalized intensity residuals for the corresponding models in (a). The progressive sharpening of the distributions around zero demonstrates the rapid convergence of the model.

**Note S4. Time-frequency analysis of the field-dependent system response.**

To demonstrate the spatial heterogeneity of the LOT system response, we analyzed the raw optoacoustic signals from three distinct regions within the imaging volume, corresponding to the exemplary PSFs shown in Fig. 3b. Fig. S4a displays the raw time-domain ultrasound response sequences collected by eight representative transducer elements for each of these three spatial locations. The corresponding one-dimensional Fourier spectra are plotted in Fig. S4b.

A clear heterogeneity is evident in both the time-domain waveforms and the position-dependent frequency signatures. For instance, the spectral content varies significantly between the three locations, indicating a spatially non-uniform system response. This variance in the raw time-frequency data, which originates from the source's position, directly translates into distinct PSF morphologies upon reconstruction using the singular value decomposition and filtered back-projection algorithms (Fig. S4c). This observation confirms the spatially variant nature of the system response and highlights the necessity of field-dependent ePSF modeling to ensure high-precision localization across the entire field-of-view (FOV).

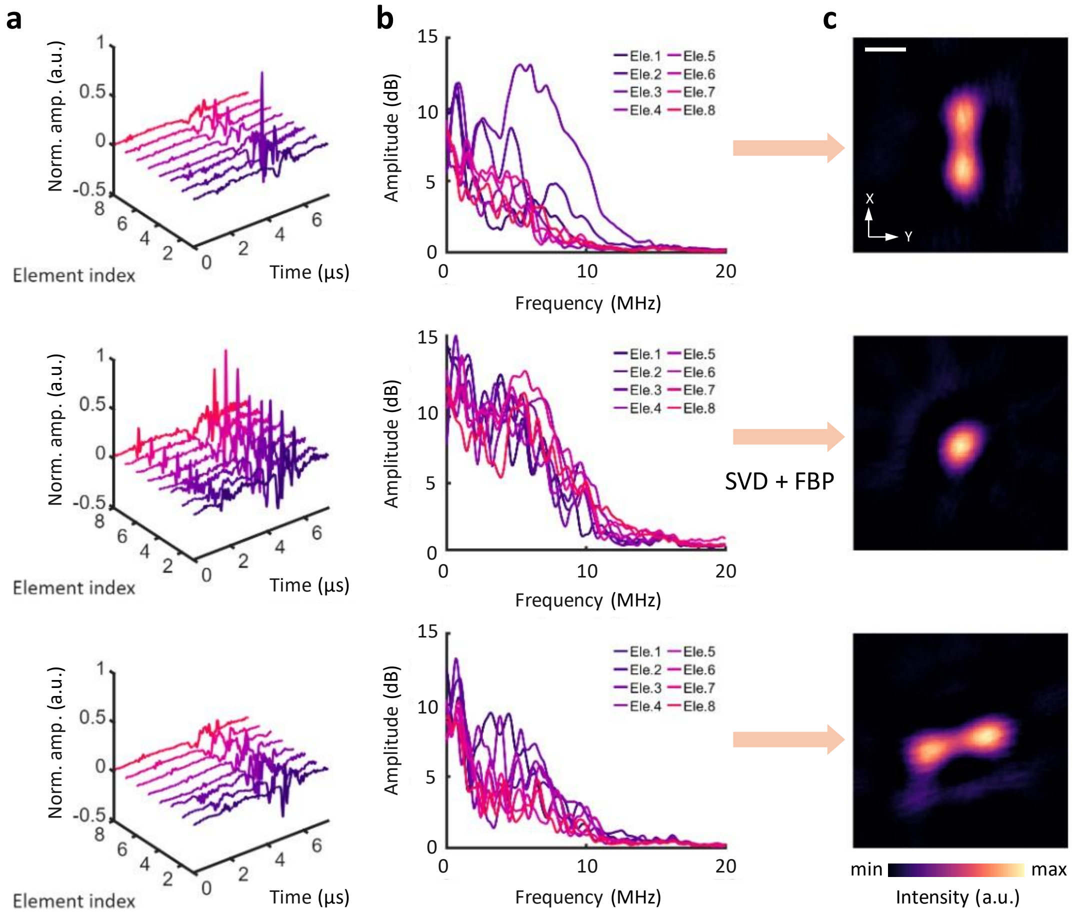


**Figure S4. Analysis of spatially variant optoacoustic signals.**

**(a)** Raw time-domain ultrasound signals from eight representative transducer elements, recorded from three distinct spatial locations (corresponding to the PSFs shown in Fig. 3b), a.u. – arbitrary units. **(b)** One-dimensional Fourier spectra of the time-domain data in (a), revealing position-dependent frequency signatures for each location. **(c)** Axial (x-y) projections of the reconstructed model generated from the raw data in (a) using singular value decomposition (SVD)-based clutter suppression and the filtered back-projection (FBP) algorithm. The clear differences in the raw signals, spectra, and final ePSF morphologies confirm the spatial heterogeneity of the system response. Scalebar – 200 µm.

**Note S5. Validation of DEPR robustness against varying noise conditions.**

To validate the effectiveness and applicability of the DEPR method for *in vivo* imaging, we evaluated its robustness against different levels of background noise. We simulated point source datasets with four distinct signal-to-background ratios (SBRs): 50:1, 20:1, 10:1, and 5:1. Using these datasets, we reconstructed the corresponding ePSF models (Fig. S5a) with a 1/4 pixels binning interval and 3 rounds of iteration.

As shown in the axial and sagittal projections, the DEPR method successfully reconstructed the ePSF model even under high-noise conditions (SBR=5:1), although the resulting model exhibited a higher background noise level compared to the high-SBR reconstructions.

We then assessed the localization precision of these reconstructed models, where point source datasets with known, randomly distributed sub-pixel positions was generated under specified SBR. Each of the four ePSF models from Fig. S5a was then used to localize these targets using the least-squares fitting (LSF) algorithm. The resulting localized results and bias distributions are plotted in Fig. S5b. Notably, even when using the ePSF model derived from the noisiest data (SBR=5:1), the average localization bias remained below 0.05 pixels. The accuracy showed only a slight degradation compared to localizations performed with the high-SBR model. The simulation confirms the robustness of the DEPR method against varying noise levels and its practical usability in realistic *in vivo* environments where SBR can be a limiting factor.

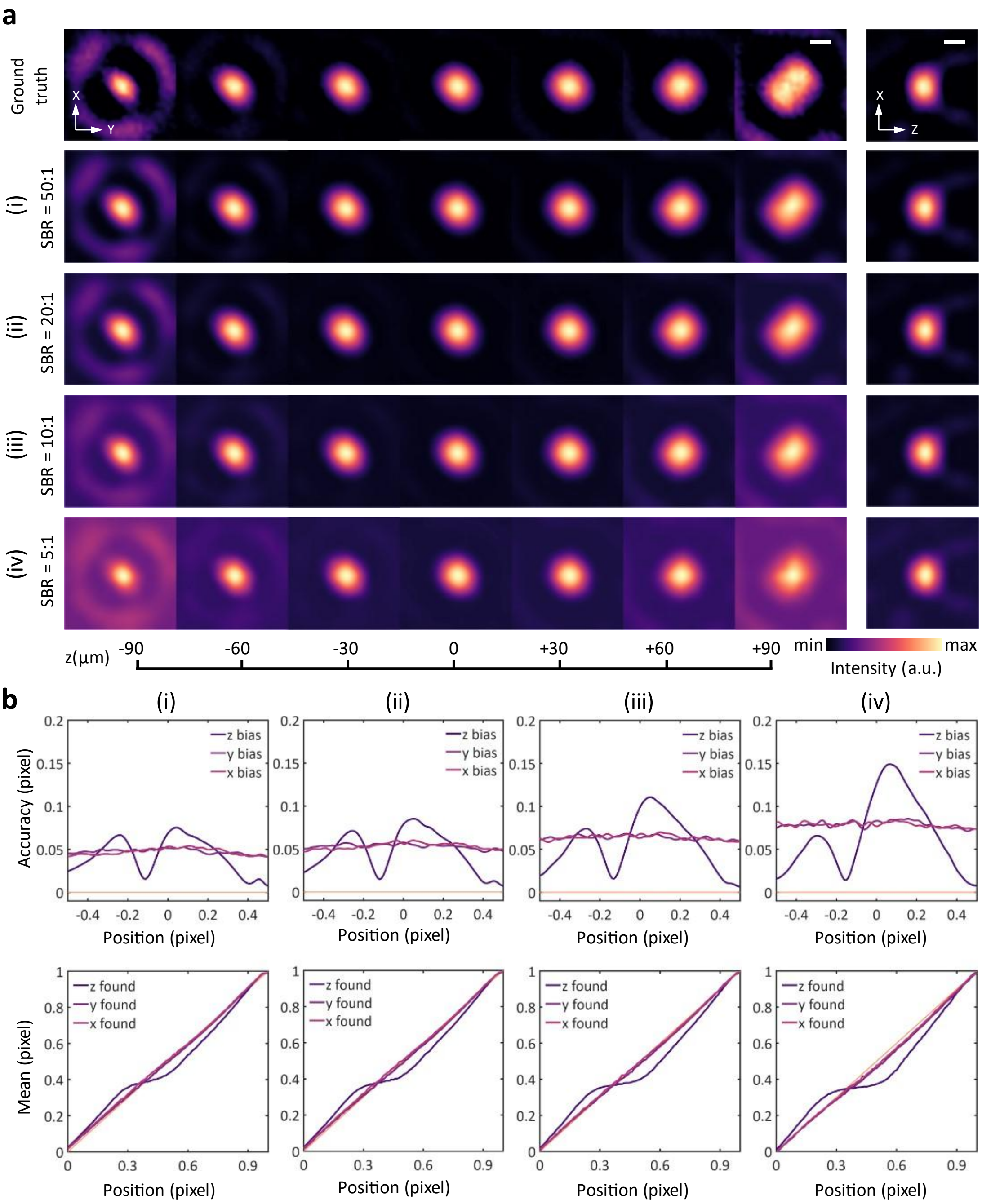


**Figure S5. Robustness of DEPR under varying signal-to-background ratio (SBR) conditions.**

**(a)** Axial (x-y) and sagittal (x-z) views of the ground truth ePSF model alongside ePSFs reconstructed from simulated datasets with SBRs of 50:1 (i), 20:1 (ii), 10:1 (iii), and 5:1 (iv). Scalebar – 100 μm, a.u. – arbitrary units. **(b)** Localization performance of the least-squares fitting (LSF) method using the corresponding ePSF models from (a). The top row plots the localization bias versus true sub-pixel position. The bottom row plots the mean localized position versus true position. The results demonstrate that high accuracy is maintained even when using the ePSF reconstructed from relatively low SBR (5:1) data.

**Note S6. Impact of the number of targets on reconstruction fidelity.**

The impact of the total number of target observations on the fidelity of the DEPR-reconstructed ePSF was further evaluated. Models were generated from simulated datasets containing 200, 300, 500, 1000, 1,500, and 2000 targets. All reconstructions used a 1/4 pixels binning interval and 3 rounds of iteration.

Qualitative results (Fig. S6a) show that with fewer than 1000 targets, the reconstructed ePSF models exhibit clear geometric distortions, as the limited number of observations is insufficient to stably recover the local structural features. At 1000 targets, these distortions are resolved, and increasing the number of targets further to 2000 only yields marginal gains. This saturation is confirmed by quantitative fidelity metrics (Fig. S6b and c). The Structural Similarity Index Measure (SSIM) between the reconstructed and ground truth models increases[6], while the Poisson deviance decreases, with both metrics showing an inflection point and plateauing around the 1000-target mark.

These findings consistently indicate that the reconstruction fidelity converges as the number of observations increases. For this simulation setup, 1000–2000 target observations provide an optimal balance between reconstruction accuracy and data acquisition efficiency.

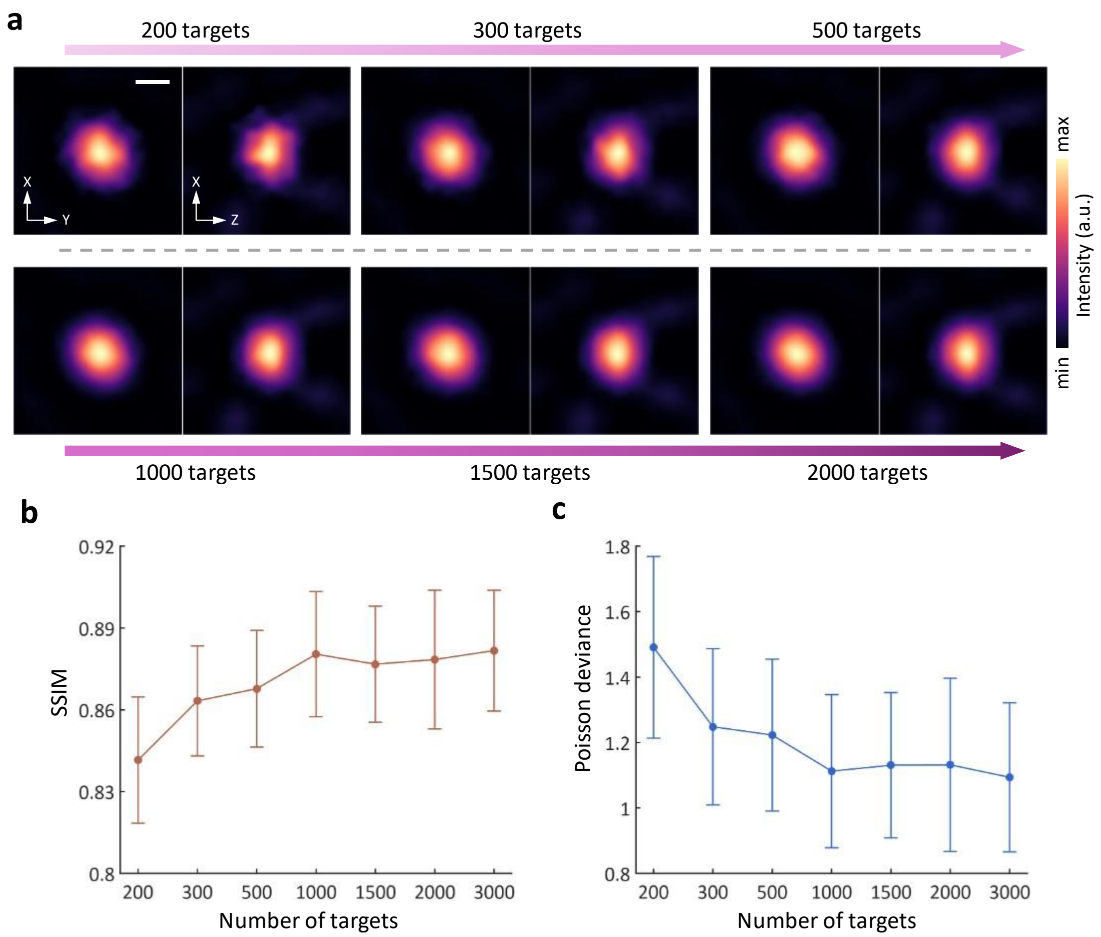


**Figure S6. Reconstruction accuracy as a function of the number of targets.**
**(a)** Axial (x-y) and sagittal (x-z) views of ePSFs reconstructed by DEPR using an increasing number of targets (200, 300, 500, 1000, 1500, and 2000). The sampling interval was set to 1/4 pixels, and 3 iterations were performed. Scalebar – 100 µm, a.u. – arbitrary units. **(b)** Structural Similarity Index Measure (SSIM) of the reconstructed ePSFs relative to the ground truth, plotted as a function of the number of targets. **(c)** Poisson deviance information metric for the reconstructed ePSFs, plotted as a function of the number of targets. For (b) and (c), data are presented as mean ± standard deviation.

**Note S7. Schematic diagram of the baseline and the modified localization workflow.**

To elucidate the differences between the localization strategies evaluated in this study, we detail the three distinct workflows illustrated in Fig. S7[7, 8]. All three commence with identical preprocessing of the reconstructed volumetric images, which involves the 'background removal' and 'initial target searching' step to identify potential candidates. The fundamental distinctions between the approaches arise in the subsequent steps of candidate detection and sub-pixel localization.

The conventional strategy, termed MT-CG (morphological thresholding - center-of-gravity), applies a 'morphological thresholding' step to filter out targets based on size or intensity. This process is intended to exclude dim or small targets that may be difficult to localize accurately, but it significantly reduces the number of targets used for reconstruction, thus lowering data utilization efficiency. Following this filtering, CG is used to determine the sub-pixel position. As a baseline for comparison, we also implemented an RM-CG (regional-maxima - center-of-gravity) approach. This method replaces the aggressive thresholding with 'regional-maxima detection', preserving a much larger number of candidates. However, it still relies on 'centroid calculation' for localization and, as such, remains highly susceptible to the pixelation-induced biases inherent to CG methods.

Our proposed DEPR-based strategy, RM-LSF (regional-maxima - least-squares fitting), also employs 'regional-maxima detection' to maximize the number of preserved targets. The critical distinction lies in the localization step. Instead of centroiding, RM-LSF utilizes LSF to match the intensity profile of each detected target against the continuous, field-dependent ePSFs retrieved by the DEPR framework. By leveraging this accurate, *in situ* system model, the RM-LSF approach robustly corrects for both pixelation effects and field-dependent aberrations. This enables highly accurate sub-pixel localization and substantially suppresses the grid-like artifacts that plague the CG-based methods.

Following the specific localization step of each pathway, all computed target positions undergo 'post-processing filtering' to remove outliers before being accumulated to render the final super-resolved LOT image.

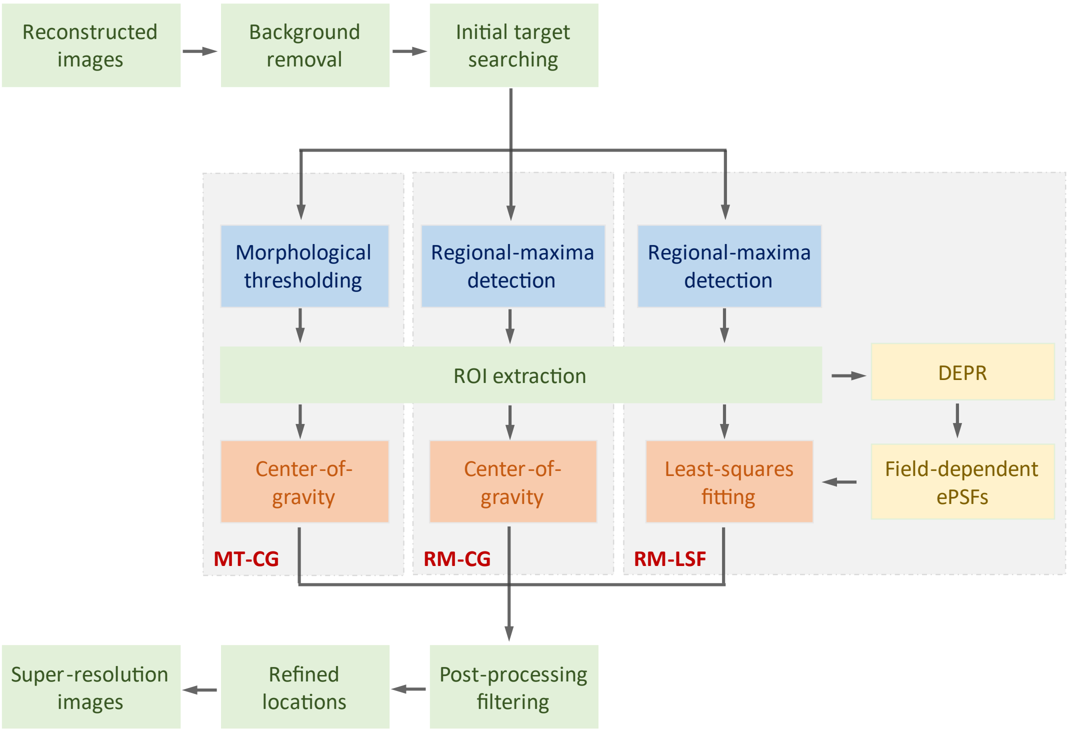


**Figure S7. Workflow comparison of localization strategies.**

Flowchart illustrating three approaches for particle localization in LOT. The conventional MT-CG (morphological thresholding - center-of-gravity) method applies morphological thresholding to exclude small targets, followed by centroid estimation. The RM-CG (regional-maxima - center-of-gravity) method removes the thresholding step and instead applies centroid estimation directly to regional-maxima detections. In contrast, the proposed DEPR-based RM-LSF (regional-maxima - least-squares fitting) method integrates *in situ* retrieved, field-dependent ePSFs with least-squares fitting, enabling precise sub-pixel localization while preserving a higher number of detected targets.

**Note S8. *In vivo* characterization of field-dependent ePSFs.**

To confirm the ability of DEPR to capture spatially variant system responses *in vivo*, we retrieved and analyzed ePSFs from the mouse brain dataset acquired under near-infrared (NIR)-I excitation with dichloromethane (DCM) microparticles. Fig. S8 displays representative ePSFs retrieved from different subregions within the imaging field. We extracted models from three adjacent regions along the horizontal (lateral) direction (upper row, (i)-(iii)) and three adjacent regions along the vertical (axial) direction (bottom row, (i)-(iii)). The line profiles along the x, y, and z axes reveal systematic differences in ePSF shape, width, and anisotropy. Notably, the horizontal profiles (top row) show clear, regular changes in the X (purple) and Y (orange) responses across the subregions. Similarly, the vertical profiles (bottom row) exhibit corresponding changes in the Z (blue) response. These observations, which are consistent with physical imaging principles[9, 10], confirm the presence of a spatially heterogeneous system response within the *in vivo* imaging volume. This analysis underscores the necessity of the field-dependent modeling provided by DEPR to ensure high-precision localization across the entire field of view.

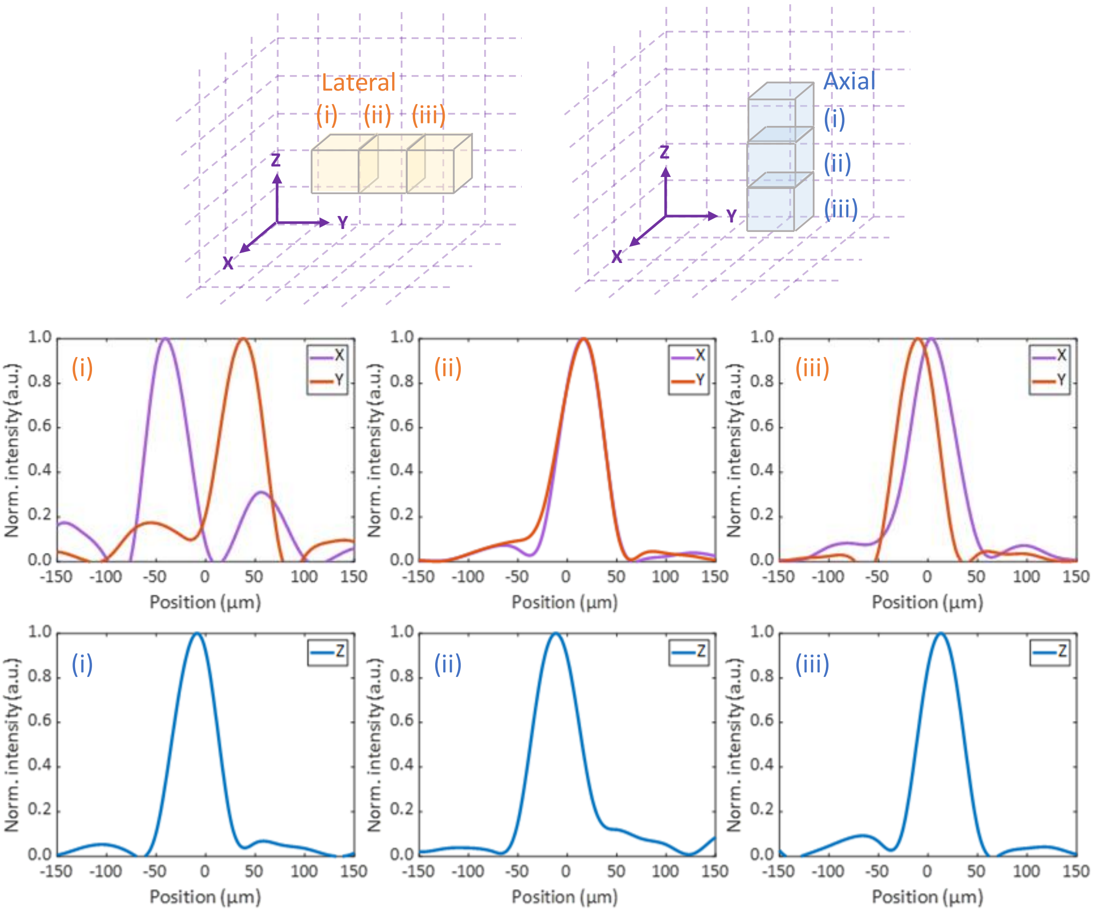


**Figure S8. *In vivo* characterization of field-dependent ePSFs.**

Representative ePSFs retrieved from different subregions of the *in vivo* mouse brain dataset acquired under near-infrared (NIR)-I excitation with dichloromethane (DCM) microparticles. Top: Corresponding X and Y line profiles for three adjacent lateral subregions (i-iii). Bottom: Corresponding Z line profiles for three adjacent axial subregions (i-iii). The systematic variations in profile shape, width, and anisotropy across the different subregions confirm the presence of a spatially heterogeneous system response *in vivo*, a.u. – arbitrary units.

**Note S9. Comparison of the rendered LOT image with respect to cumulative frames.**

To assess the image enhancement and compare the performance of the three localization strategies (MT-CG, RM-CG, and RM-LSF), we reconstructed the *in vivo* mouse brain vasculature acquired under NIR-II excitation with $CaCO_3$@CuS/PDA microparticles using cumulative data. Axial (x-y) and sagittal (y-z) projections were generated at 4000, 8000, 12000, and 16000 frames.

As shown in Fig. S9, the structural density and overall signal intensity improved for all three methods as the number of accumulated frames increased. However, significant performance variations were evident. The conventional MT-CG strategy (top row) consistently produced reconstructions with sparse vascular networks and lower signal intensity. This is a direct consequence of the morphological thresholding step, which discards a large fraction of valid target signals and thus suffers from low data utilization efficiency.

In contrast, the RM-CG method (middle row), which omits this thresholding step, captured a much higher density of localizations. While this improved structural completeness, its reliance on conventional CG approach introduced severe, grid-like artifacts that distort the image and obscure fine details, particularly visible in the sagittal projections.

The proposed RM-LSF strategy (bottom row) demonstrated superior performance across all frame counts. It successfully combined the high data utilization of regional-maxima detection with the high accuracy of DEPR-based localization. Consequently, the RM-LSF reconstructions consistently revealed the finest structural details, with enhanced contrast and a significant reduction in the gridded artifacts that plagued the RM-CG method. This progressive comparison confirms that RM-LSF provides the most faithful and detailed representation of the microvasculature.

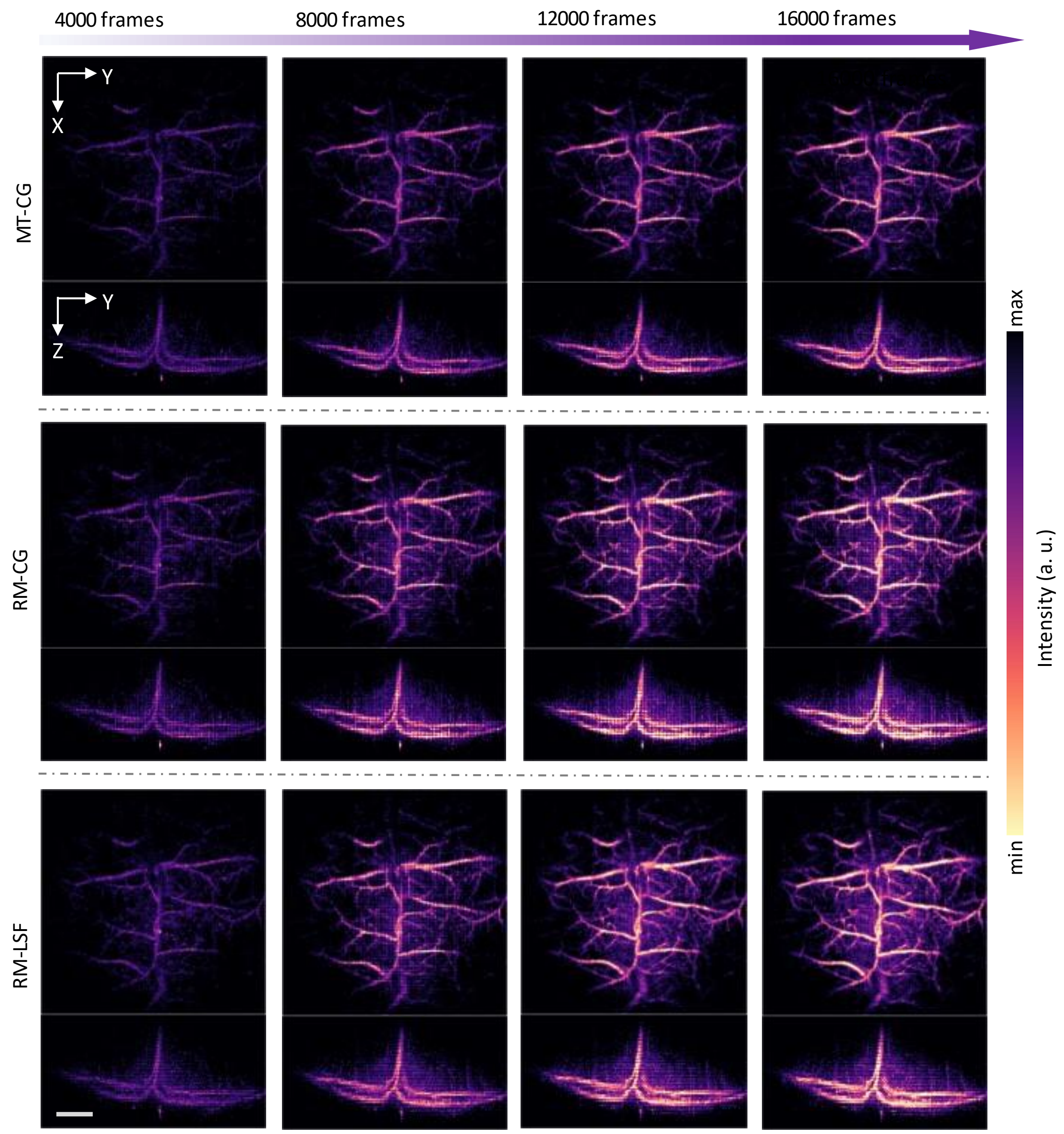


**Figure S9. Progressive enhancement of LOT image quality with cumulative frames.** Comparison of the three localization strategies using the *in vivo* NIR-II dataset, reconstructed with 4000, 8000, 12000, and 16000 cumulative frames. The rows correspond to the MT-CG (top), RM-CG (middle), and RM-LSF (bottom) methods. Each panel displays the axial (x-y, top) and sagittal (y-z, bottom) projections. The RM-LSF method consistently reveals superior structural detail and reduced gridded artifacts, outperforming the other two methods. Color intensity represents accumulated particle density. Scalebar – 1 mm, a.u. – arbitrary units.

**Note S10. Computational performance comparison of localization methods.**

We compared the computational performance of the DEPR-based LSF localization method against the conventional CG method. The analysis was performed by measuring the processing time required for an increasing number of localized points (from 10000 to 90000), with results averaged over 50 independent runs.

As shown in Fig. S10, the total computation time for both methods scales linearly with the number of points processed, as indicated by the dashed trend lines. The LSF method (orange) consistently exhibits a slightly higher computational cost than the CG method (blue). This modest overhead is expected due to the iterative optimization nature of LSF, whereas CG is a non-iterative calculation.

The average time per point remains relatively constant regardless of the total number of points processed. On average, LSF required approximately 12.5 µs per point, compared to 10.5 µs per point for CG, which represents a minor computational overhead. Given the significant improvements in localization accuracy and the substantial suppression of gridded artifacts provided by LSF, this modest increase in computation is well-justified. Furthermore, the total processing time for the entire super-resolution pipeline remains dominated by the initial image reconstruction, not the localization step.

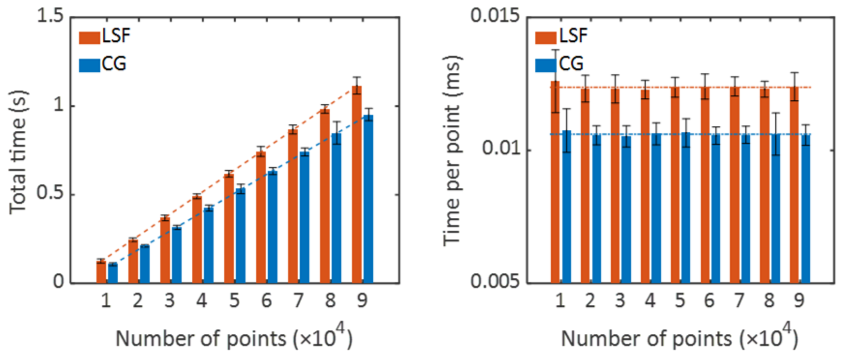


**Figure S10. Computational performance comparison of localization methods.**
**Left:** Total computation time versus the number of localized points for LSF (orange) and CG (blue). Both methods demonstrate linear scaling. **Right:** Average time per point, which remains relatively constant. LSF exhibits a modest computational overhead compared to CG. All results are averaged over 50 independent runs, error bars represent standard deviation.
